# KDM3A and KDM3B Maintain Naïve Pluripotency Through the Regulation of Alternative Splicing

**DOI:** 10.1101/2023.05.31.543088

**Authors:** Caleb M. Dillingham, Harshini Cormaty, Ellen C. Morgan, Andrew I. Tak, Dakarai E. Esgdaille, Paul L. Boutz, Rupa Sridharan

## Abstract

Histone modifying enzymes play a central role in maintaining cell identity by establishing a conducive chromatin environment for lineage specific transcription factor activity. Pluripotent embryonic stem cell (ESC) identity is characterized by a lower abundance of gene repression associated histone modifications that enables rapid response to differentiation cues. The KDM3 family of histone demethylases removes the repressive histone H3 lysine 9 dimethylation (H3K9me2). Here we uncover a surprising role for the KDM3 proteins in the maintenance of the pluripotent state through post-transcriptional regulation. We find that KDM3A and KDM3B interact with RNA processing factors such as EFTUD2 and PRMT5. Acute selective degradation of the endogenous KDM3A and KDM3B proteins resulted in altered splicing independent of H3K9me2 status or catalytic activity. These splicing changes partially resemble the splicing pattern of the more blastocyst-like ground state of pluripotency and occurred in important chromatin and transcription factors such as *Dnmt3b, Tbx3* and *Tcf12*. Our findings reveal non-canonical roles of histone demethylating enzymes in splicing to regulate cell identity.

## INTRODUCTION

Gene expression patterns are established by the coordination of transcription factor binding to chromatin (1–3). The chromatin environment is controlled in part by chemical modifications on histones and DNA (4, 5). In general, histone acetylation decompacts chromatin to facilitate transcription factor binding and promotes gene expression (6, 7). The effect of histone methylation is more nuanced and depends on the specific amino acid residue that is modified (8, 9). The abundance of histone modifications at a particular location is controlled by the balance of the activities of enzymes that add or remove each modification that is modulated by the protein complexes they associate with (10, 11). As part of such protein complexes, histone modifying enzymes may serve scaffolding and targeting functions independent of their catalytic activity. Hence histone modifying enzymes with the same catalytic targets can have both redundant and distinct functions.

Histone 3 Lysine 9 dimethylation (H3K9me2) is correlated with gene repression and enriched in tightly compacted heterochromatin (12, 13). H3K9me2 is found on the majority of histone tails in somatic cells and enriched in broad megabase domains that encompass both repetitive elements and protein coding genes (14–17). The enzymes KDM3A and KDM3B demethylate H3K9me1/me2 and have conserved catalytic domains (18–21). In mice, the combined knockout of both KDM3A and KDM3B leads to embryonic lethality at day 4.5 when the epiblast is being formed, insinuating the redundant function of the two enzymes (22). In contrast, while the KDM3A mouse knockout is obese but viable (23–25), the KDM3B mouse knockout is perinatal lethal, with one-third of the offspring dying within 48 hours of birth (26, 27).

KDM3A and KDM3B also have distinct functions in gaining pluripotency from somatic cells but overlapping functions in the maintenance of pluripotency. Somatic cells can be reprogrammed into induced pluripotent stem cells (iPSCs) by the overexpression of the Yamanaka transcription factors (28). In this transition, the knockdown of *Kdm3b* severely compromises reprogramming efficiency although KDM3A levels remain unaltered (29, 30). In contrast, the self-renewal of pluripotent embryonic stem cells ESCs is compromised only upon the deletion of both KDM3A and KDM3B (22). Interestingly the KDM3A/KDM3B double knockout resulted in few gene expression changes which is unexpected from a broadly repressive modification (22).

To investigate how KDM3A and KDM3B functions are modulated by their protein interactions, we performed immunoprecipitation of KDM3A and KDM3B in ESCs. We found both enzymes interact with the RNA splicing machinery. By performing acute degradation of both proteins using a “degron” tag for targeted ubiquitination and proteolysis (31, 32), we found deficits in splicing. These splicing changes were independent of H3K9me2 status of the associated exons and could be rescued by catalytically inactive versions of the KDM3 proteins. The splicing aberrations occurred in several genes that are important regulators of cell state transitions, mimicking the splicing patterns of a more blastocyst like state of pluripotency. Taken together we uncover a role for KDM3A and KDM3B in splicing regulation that controls cell identity during development.

## MATERIALS AND METHODS

### Cell culture

Tet-inducible V6.5 ESC lines (33) and derivatives were routinely maintained on 0.1% gelatin coated plates in ESC media (KnockOut DMEM, 15% fetal bovine serum (FBS), Glutamax, Penicillin/Streptomycin, non-essential amino acids (NEAA), 2-mercaptoethanol (4 μL/500 mL), and leukemia inhibitory factor (LIF) with irradiated DR4 feeder mouse embryonic fibroblasts (MEFs). E14 ESCs and derivatives grown in ESC media in feeder-free conditions on 0.1% gelatin coated plates; all ESCs confirmed to be mycoplasma free. For growth in 2i/LIF conditions, ESCs were maintained on 0.1% gelatin coated plates in modified 2i/LIF media: 50% DMEM/F12 (Gibco), N-2 Supplement (17502048), Insulin (12.5 mg/L0), progesterone (0.01 mg/L), 50% Neurobasal medium (Gibco), B-27 minus Vitamin A (Thermo Fisher 12587010), 1000 U/mL LIF, b-mercaptoethanol (6.3 ul/L), Penicillin-Streptomycin (Gibco), 3 µM CHIR-99021 (Reprocell 04-0004), 0.4 PD-0325901 (Reprocell 04-0006), Bovine Serum Albumin (0.0025%), and 5% Fetal Bovine Serum. Passaged every 2-3 days with TrypleE (Gibco).

### Cell line derivation

3xFLAG-KDM3A and 3xFLAG-KDM3B Tet-inducible V6.5 ESC lines were generated by FLP-mediated recombination of N-terminal 3X-FLAG-tagged KDM3A or KDM3B at the Col1A1 locus (33). Cells were selected with hygromycin at a final concentration of 100 µg/ml and plated at low density to isolate single clones and screened via Western Blot for endogenous KDM3A or KDM3B and FLAG (M2 FLAG Sigma). FLAG-tagged protein expression was induced with 1ug/ml doxycycline.

Degron cell lines were generated following (31, 32). Briefly, *Kdm3a* microhomology arms were added to the pCRIS-PITChv2-Puro-dTAG (Addgene #91793) donor vector and *Kdm3b* microhomology arms were cloned into the pCRIS-PITChv2-BSD-dTAG (Addgene #91792) donor vector to allow for dual selection with puromycin and blasticidin S hydrochloride.

Cas9/sgRNA targeting vectors were constructed according to Sakuma et al. 2016. Briefly, pX330A-1x2 (Addgene #58766) and pX330S-2-PITCh (Addgene #63670) were combined with guide RNAs (gRNAs) targeting *Kdm3a* (5’-GAAACCATGGTGCTCACGCT-3’) or *Kdm3b* (5’-CAACAGCCGCTTGCCCACCG-3’) to generate a vector PITCh sgRNA, target-specific sgRNA, and Cas9. Two million E14 ESCs were nucleofected with paired *Kdm3b* targeting and donor plasmids (5 µg each) using the Amaxa 4D-Nucleofector protocol (Lonza) and selected with blasticidin S hydrochloride (Research Products International, B12200) at a final concentration of 10 ug/ml. After selection, ESCs were plated at low density to isolate single clones and screened via Western Blot for KDM3B or HA. A clone expressing degradable KDM3B was selected and nucleofected through the same procedure with 5 µg of the *Kdm3a* targeting and donor plasmids, selected with puromycin at 5 ug/ml, plated at low density to isolate single clones and screened by western blot for endogenous KDM3A and HA. Clones were then expanded and normal karyotype was confirmed. All degron degradations were induced using dTAG-13 (Sigma-Aldrich SML2601) at a final concentration of 500nM.

The *Kdm3b*-KO ESC line was generated in the E14 background by using two gRNAs targeting exon 19 (5’-ACGAGATGGCAGGCTCAATC -3’) and exon 22 (5’-CTTCAAAGAGTCGCTTTCGG -3’), which were inserted into dual gRNA targeting vector pX333 (Addgene #64073). E14 ESCs were then transfected with 0.5µg of the targeting vector using Lipofectamine 3000 (Thermo Fisher Scientific-L3000001). Transfected ESCs were then plated at low density to isolate single clones and screened via immunofluorescence and Western Blot for endogenous KDM3B.

Full length and catalytic mutant KDM3A overexpressing cells were generated in WT E14 ESCs and the KDM3A/KDM3B degron ESCs through the following methods. First, the vector pCDH-FLAG was linearized with NheI and EcoRI. Kdm3a was amplified from cDNA with NheI and EcoRI overhangs and ligated together with T4 ligase. This resulted in pCDH-FLAG-Kdm3a-FL, which was then used in a quickchange PCR with mispriming primers (5’-ATCTTGGCTTAAATGTATCTGATGCAGC -3’ and 5’-ATACATTTAAGCCAAGATTTGTGGTCCC -3’). PCR product was then digested with DpnI to get rid of template and transformed. Clones were screened with sanger sequencing and confirmed to have catalytic inactive H1120G/D1122N mutations (34, 35). Next, ESC lines were generated using lentiviral transfer plasmids pCDH-Flag-Kdm3a-FL and pCDH-Flag-Kdm3a-CM. Lentiviral vectors were transfected into HEK293T cells with packaging vector psPAX2 (Addgene #12260) and envelop vector VSV-G using polyethylenimine (PEI 1mg/ml). Media was changed to ESC media with 10mM sodium butyrate and 20 mM HEPES 16 hours post-transfection. Media was then changed to ESC media with 20 mM HEPES 7 hours post-sodium butyrate induction. Virus-containing media was harvested at 48h and 72h, combined, and filtered through a 0.45 μm PVDF filter. Virus from virus containing media was then concentrated using homemade LentiX concentrator (40% PEG-8000, 1.2M NaCl) overnight at 4°C rotating. Concentrated virus was transduced into pre-plated ESCs at a 1:200 concentration with 10 μg/ml Hexadimethrine Bromide (polybrene). After selection with G418, batches were screened for overexpression using FLAG western blots.

Feeder MEFs were isolated and maintained in MEF media (DMEM, 10% fetal bovine serum (FBS), Glutamax, Penicillin/Streptomycin, non-essential amino acids (NEAA), 2-mercaptoethanol (4 μL/500 mL) from DR4 mice genetically resistant to geneticin (G418), puromycin, hygromycin, and 6-thioguanine. Feeder cells were irradiated with 9,000 rad after three passages. Mice were maintained according to the UW-Madison institutional animal care and use committee (IACUC)-approved protocol.

### RT-qPCR Analysis and semi-quantitative rMATs validation

RNA from cell pellets treated with dTAG-13 for four hours was isolated using the Isolate II RNA Mini Kit (Bioline BIO-52072), and 1 μg was converted to cDNA using the qScript cDNA Synthesis Kit (VWR 101414). Technical replicates of 20 ng (based on RNA concentration) were used to measure Ct on a BioRad CFX96 thermocycler with iTaq UniverSYBR Green SMX (BioRad 1725125) in 20 µl reactions. Relative expression was normalized to the housekeeping gene *Gapdh*.

For targets used in the KDM3A rescue experiments (Figure 5), targets from the rMATs analysis were chosen by a PSI level of at least 15%, a maximum gene expression difference of 1.7x fold TPMS and average minimum junction read count of 65 for the included isoform. Differential splicing events were analyzed by semi-quantitative PCRs, performed at 40 cycles for *Plod2* and *Lsm14b*. PCR products were observed through gel electrophoresis and band intensities were quantified using Licor software and reported as inclusion and skipped splicing ratio.

For 2i/LIF related splicing validation (Figure 6), rMATS validation primers were designed using PrimerSeq (36) to distinguish differential splicing events and semi-quantitative PCRs were performed for X cycles dependent on target (27 cycles *Dnmt3b,* 35 cycles *Tbx3,* 40 cycles *Tcf12*) and analyzed by gel electrophoresis. Gel intensity from each experiment was analyzed by Licor software and reported as arbitrary fluorescence units.

Primers used for splicing analysis:

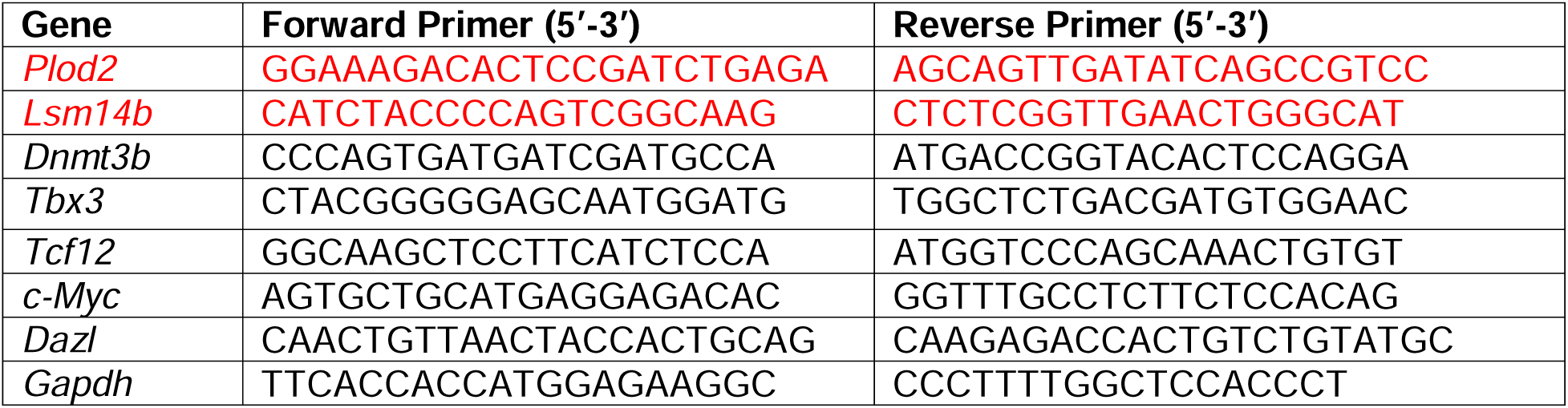

### Immunoblot

Whole cells were lysed in RIPA buffer (150 mM NaCl, 1% NP-40, 0.5% Na deoxycholate, 0.1% SDS, 25 mM Tris pH 7.4) with 1x protease inhibitor (Roche 04693116001), sonicated for 5 secs at 20% amplitude, and quantified with the DC Protein Assay (BioRad 5000112) according to the manufacturer’s instructions. Equal amounts of protein were loaded onto an SDS-Page gel and transferred to a nitrocellulose membrane. Membranes were blocked in blocking buffer (5% milk, 0.1% Tween-20, 1x PBS) followed by incubation with primary antibody in blocking buffer.

Membranes were washed 0.1% Tween-20-PBS and incubated with secondary antibody in blocking buffer. Membranes were washed and visualized with ECL reagent. Images were quantified using Image Studio Lite software. Primary antibodies: anti-H3K9me2 (Abcam ab1220, 1:1000, lot#GR33089025), anti-H3K9me3 (Active Motif 39161,1:1000, lot#01022004), anti-KDM3A (Proteintech 12835-1-AP,1:1000, lot#00065897), anti-KDM3B (U.S. Biological 1:1000, lot#L20080658), anti-EFTUD2 (LS-BIO-C346198, 1:1000, lot#207907), anti-SNRNP200 (Proteintech 23875-1-AP, 1:1000), anti-PRMT5 (EMD Millipore 07-405, 1:250, lot#3288011) anti-RNAPII (Cell Signaling 14958, 1:1000, lot#5), anti-α-TUBULIN (Cell Signaling 3873, 1:2500, lot#16), anti-FLAG M2 (Sigma-Aldrich, F3165, 1:1000).

### Immunofluorescence

Performed as previously described (37). Briefly, coverslips were fixed in 4% paraformaldehyde-PBS, permeabilized in 0.5% Trition-X-PBS, and washed in 0.2% Tween-20-PBS. Coverslips were blocked for 30 min in blocking buffer (1x PBS, 5% donkey serum, 0.2% Tween-20, and 0.2% fish skin gelatin). Cells were stained for 1 hr with primary antibody in blocking buffer, rinsed 2x in wash buffer, and stained for 1 hr with secondary (1:1000) in blocking buffer.

Coverslips were rinsed with wash buffer, stained with DAPI (0.1 μg/ml) in wash buffer, and rinsed with wash buffer. Imaging was performed on Nikon Eclipse Ti using NIS Elements software at 20x zoom. Primary antibodies: anti-HA (Santa Cruz sc7392, 1:250, lot#G0622).

### Duolink Proximity Ligation Assay

Coverslips were prepared as in the Immunofluorescence section and then stained with the Duolink® In Situ Detection Reagents Red (DUO92008) following the manufacturer’s instructions. Briefly, coverslips were blocked for 30 minutes at 37°, stained with primary antibody in Duolink antibody dilutent for 1 hr at room temperature, incubated with Duolink mouse and rabbit probes for 1 hr at 37°, ligated for 30 minutes at 37°, amplified for 100 minutes at 37°, and finally mounted with Aquapolymount (Polysciences, 18606). Confocal images were acquired using a Nikon A1R confocal microscope with Nikon NIS-Elements software and analyzed with NIS-Elements. Z-stacks were taken in 1 µM slices, approximately 5 slices taken per colony, with slice in the middle of colony selected. PLA foci were quantified in ImageJ using Image->Adjust->Threshold followed by Analyze->Analyze Particles to quantify individual interactions. Nuclei were then quantified using the ImageJ cell counter tool and the ratio of particle number to number of nuclei calculated. For the transcriptional inhibition PLA, cells were treated with 5,6-Dichloro-1-beta-Ribo-furanosyl Benzimidazole (DRB) for 3 hrs at a final concentration of 10µM and then processed at the same Duolink™ parameters above and finally imaged with Nikon Eclipse Ti using NIS Elements software. Primary antibodies: anti-HA (Santa Cruz sc7392, 1:250, lot#G0622), anti-PRMT5 (EMD Millipore 07-405, 1:250, lot#3288011), anti-KDM3B (Cell Signaling 5377, 1:250, lot#1), anti-EFTUD2 (LS-BIO-C346198, 1:250, lot#207907), and anti-OCT4 (Cell Signaling 2840, 1:250, lot#16).

### Protein Extraction and Immunoprecipitation

Total protein was extracted using a modified nuclear extraction method (38) as previously described (39). Briefly, ESCs were feeder depleted, and extracted in 5x volume buffer A (10 mM HEPES [pH 7.9],1.5 mM MgCl2, 10 mM KCl, 1 mM DTT, Phosstop™-phosphatase inhibitor (Roche 04906837001) and cOmplete™ EDTA-free Protease Inhibitor Cocktail (Roche 04693132001) for 15 minutes on ice after gentle douncing. Nuclei were pelleted at 1500xg for 10 minutes, washed in buffer A, and resuspended in 2x volume buffer C (0.42 M KCl, 20 mM HEPES [pH 7.9], 0.2 mM EDTA [pH 8.0], 5% glycerol, 1 mM DTT, and phosphatase and protease inhibitor cocktail and dounced thoroughly. After douncing, MgCl_2_ was spiked into a final concentration of 1.5 mM, benzonase added (250 units/ml; Sigma-Aldrich E1014), and incubated for 10 minutes at room temperature. Extracts were then rotated at 4°C for 2 hours. Soluble nuclear extracts were isolated by centrifugation at 23,000xg for 30 minutes at 4°C.

For FLAG-IP-MS, 10-15 mg of nuclear extract was dialyzed overnight to 0.3M KCl using a 7,000 MWCO Slide-A-Lyzer Dialysis Cassette G2 (Thermo Scientific). Extracts were then incubated with FLAG-M2 agarose (Sigma, 50% slurry) bead slurry at a ratio of 90 ul beads slurry to 400 ug of dialyzed protein for 2 hours rotating at 4°C. Samples were washed thrice with IP buffer (0.3 M KCl, 20 mM HEPES, and phosphatase and protease inhibitor cocktail) and collected by spinning at 1000g x minutes between washes. IP complexes were eluted with 200 ng/uL of 5X FLAG peptide (manufactured by Pepmic).

Eluted protein complexes were trichloroacetic acid (TCA) precipitated, in-solution digested with trypsin (40), desalted using Agilent Bond Elut OMIX C18 SPE pipette tips per manufacturer protocol and eluted in 10µl of 60/40/0.1% ACN/H_2_O/TFA. Dried to completion in the speed-vac and finally reconstituted in 40µl of 0.1% formic acid. Peptides were analyzed by nanoLC-MS/MS using the Agilent 1100 nanoflow system (Agilent) connected to hybrid linear ion trap-orbitrap mass spectrometer (LTQ-Orbitrap Elite™, Thermo Fisher Scientific) equipped with an EASY-Spray™ electrospray source (held at constant 35°C). Chromatography of peptides prior to mass spectral analysis was accomplished using capillary emitter column (PepMap® C18, 3µM, 100Å, 150x0.075mm, Thermo Fisher Scientific). Protein annotations, significance of identification and spectral based quantification was done with Scaffold software (version 4.11.0, Proteome Software Inc., Portland, OR). Peptide identifications were accepted if they could be established at greater than 7.0% probability to achieve an FDR less than 1.0% by the Scaffold Local FDR algorithm. Protein identifications were accepted if they could be established at greater than 99.0% probability to achieve an FDR less than 1.0% and contained at least 2 identified peptides.

For HA-Tag Co-IP, 1 mg of nuclear extract of KDM3A and KDM3B endogenously tagged ESCs (prepared as above) was incubated with 10 µg of HA antibody (Roche, clone 12ca5, catalog#11583816001) for 4 hours rotating at 4°C. Immunocomplexes were collected with 50ul of Protein G Dynabeads (Invitrogen, catalog#10003D) by rotation at 4°C for one hour. Samples were washed thrice with IP buffer (0.3 M KCl, 20 mM HEPES, and phosphatase and protease inhibitor cocktail), and collected by magnet. IP complexes were sequentially eluted first with 50 ul of 1 mg/ml HA peptide (Sino Biological, catalog #PP100028, lot #IP15MA1401X) then boiling in sample buffer 95°C for 10 minutes.

### Crapome Filtering and String Network Analysis

KDM3A and KDM3B FLAG-IP mediated interactome was first filtered for non-nuclear compartments using DAVID GO analysis (41, 42) (https://david.ncifcrf.gov/), then remaining proteins were further filtered using the “crapome” contaminant repository (https://reprint-apms.org/)(43). Contaminant controls (cc2,cc3,cc4,cc5,cc6,cc28) were selected based on similarity of nuclear extraction method, instrumentation, and sensitivity along with a FLAG-IP from our uninduced ESCs. After analysis with the Crapome pipeline, proteins with a minimum spectral count of 3, saint probability score of 0.7, and fold change (FC) cutoff of 2 were kept for further analysis. String network analysis (https://string-db.org/) was performed on the overlapping or unique interactomes of KDM3A and KDM3B. Experiments and databases were used for active interaction sources at a high confidence, hiding disconnected nodes. The resulting network was exported to Cytoscape (44) for further visual editing based on FC. The largest bubbles have a FC >10, medium FC 10 <x> 5, and small FC x<5.

### RNA isolation and library preparation

Samples for RNA-seq were prepared as previously described (45). ESCs (WT and degron) were grown in three independent replicate samples and treated with dTAG-13 for 4 hours, washed with PBS, harvested with trypsin and stored in Trizol (Sigma 15596018). RNA was isolated by adding one-fifth the volume of chloroform to each sample and phases separated by max centrifugation. The upper aqueous layer was isolated, RNA was precipitated with 0.53 volumes of ethanol, and applied to a RNeasy column (Qiagen 74104). RNA was washed with 500 μl RW1 and DNA was digested on the column with DNase (Qiagen 79254) for 30 min. RNA was washed according to the RNeasy protocol and eluted in 30 ul of H2O. RNA was quantitated with Qubit, and 1 μg of RNA was used as input for library preparation. The cDNA library was constructed using TruSeq RNA Sample Preparation kit V2 (RS-122-2002) according to the manufacturer’s instructions. Libraries were assessed with Bioanalyzer3.0.

### RNA-Seq computational and statistical analysis

RNA-seq analysis was performed as described in (46). Libraries were sequenced to a depth of ∼50 million reads/sample were sequenced PE150 by Novogene on a Illumina Novaseq platform. Sequencing quality was assessed with FastQC (http://www.bioinformatics.babraham.ac.uk/projects/fastqc/). Sequenced reads were processed using Trimmomatic (47) with the following parameters: LEADING:3 TRAILING:3 CROP:135 HEADCROP:15 SLIDINGWINDOW:4:15 MINLEN:36. Reads were aligned to the mm9 genome using RSEM-1.2.4 (48) with a mismatch per seed set to 2 and seed length set to 28 (-bowtie-m 200 --bowtie-n 2 --forward-prob 0.5 --seed-length 28 --paired-end). RSEM-1.2.4 alignment yielded transcripts per million (TPM) for each gene. A median TPM value from the three replicates was used for all main figures. Differentially expressed (DE) genes were called with EBSeq (49). Differentially expressed genes were filtered to have a posterior probability DE greater than 0.95 and a 2 or 0.5 greater posterior fold change (PostFC). Expression changes reported are PostFC values determined by EBSeq unless otherwise noted to be TPM. Gene Ontology was performed using DAVID (41, 42). DAVID parameters included: GOTERM_BP_DIRECT or BP_5 and GOTERM_MF_DIRECT or MF_5, viewed by functional annotation clustering. GO terms within functional annotation clusters with a p-value X<.001 were considered significant.

### Differential Splicing Analysis

Sequencing reads were first mapped to the mouse genome, mm9, using STAR aligner (version 2.7.8a) (50). The mm9 genomic index was created using the genomegenerate function from STAR with ENCODE fasta and comprehensive gene annotation (NCBIm37). To identify canonical differential splicing events, mapped data was analyzed with rMATS 4.1.2 (51–53) was used for counting junction reads amongst three replicates of each condition. Filtering for significant splicing events was based on FDR<0.1, minimum inclusion level difference of 5%, and 15 minimum read count in at least one included or skipped isoform. Heatmap generated using pheatmap in Rstudio with kmeans clustering of 5 for inclusion level difference of aberrantly spliced genes.

Transcriptomic annotation was performed as originally described (54) with some updates. Mapped reads were filtered to remove repeat RNAs and exonic reads. Reads attributable to alternative polyadenylation sites were removed based on 3’ end sequencing of mouse embryonic stem cells. Intronic reads were partitioned and allocated to each intron proportional to its weight assuming all introns of a gene are present in equal abundance. Differential analysis using DESeq was then used to determine introns enriched in read coverage (observed read counts) compared to the expected counts using an FDR adjusted p-value threshold of 0.01 and fold change threshold of 2. We used custom scripts to derive a transcriptome annotation based on Mm9 gene start and end boundaries, the genomic locations of detained introns as determined above, and the genomic locations of all splice junctions. To identify intronic polyadenylation (alternative last exon (ALE) choice) events and detained introns, the Dexseq (55) pipeline was used. Events were counted using the dexseq_count.py script of mapped reads against a custom annotation for mouse ESCs previously generated in (54). For the data presented in Figure 4, only events with a padj value<0.1 and from events annotated as Detained Introns or Class I ALEs were used. All scripts are available on request.

### ChIP-qPCR analysis

Methodology for chromatin immunoprecipitation (ChIP)-qPCR was adapted from ChIP-seq procedures as previously described (56). 2×15 cm of WT E14 ESCs or KDM3A/KDM3B degron ESCs (Uninduced or +dTAG-13 for 4 hours) were used as starting material. ChIP was performed on a separate biological replicate expansion, and duplicates were immunoprecipitated on separate days. Cells were trypsinized and fixed with 1% formaldehyde in suspension for 10 min, rotating. Cross-linking was quenched with 0.14 M glycine for 5 min, cells were centrifuged at 300g for 3 min, and pellets were washed three times with cold 1× PBS. Cells (12.5 million) were resuspended in 0.100 ml of lysis buffer [1% SDS, 50 mM tris-HCl (pH 8.0), 20 mM EDTA, 1× cOmplete (Roche, 4693132001) protease inhibitor] and sonicated on a Covaris S220 Focused-ultrasonicator with the following parameters:15 cycles of 45 seconds ON (peak 170, duty factor 5, cycles/burst 200), 45 s OFF (rest) in 6° to 8°C degassed water. Aliquot (10 μl) taken at 15 cycles to check DNA fragmentation. Aliquots were diluted with 75 μl of H2O and incubated with 10 μg of ribonuclease (RNase) for 10 min at 37°C. Two microliter of proteinase K (10 mg/μl) was added, and sonication samples were incubated at 60°C overnight to reverse crosslinks. DNA was purified with phenol-chloroform extraction with phase lock tubes, precipitated with ethanol, and run on a 1.5% agarose gel to ensure generation of 200- to 400-bp fragments. Sonicated chromatin was centrifuged at 21,000*g* at 6°C for 10 min, and supernatant was collected and quantified with the Qubit DNA HS Assay Kit (Thermo Fisher Scientific, Q32854). Chromatin was aliquoted ensuring that the SDS remained in solution so that the concentration was exactly 0.1% after diluting 1:10 in dilution buffer [16.7 mM tris-HCl (pH 8.0), 0.01% SDS, 1.1% Triton-X, 1.2 mM EDTA, and 167 mM NaCl].

H3K9me2 and H3K36me3 were immunoprecipitated with 5 ug of the following antibodies: H3K9me2 (Abcam, #1220) and H3K36me3 (Active Motif, #61902) from 10 μg of chromatin. ChIPs were incubated overnight at 4°C with rotation, and then Dynabeads were added for 2 hours. Dynabeads were pre-prepared as follows: 25 μl of protein A (Thermo Fisher Scientific, 10002D) and 25 μl of protein G (Thermo Fisher Scientific, 10004D) were combined and washed once in PBS, 0.02% Tween 20 and once in dilution buffer, and then resuspended in an equal volume of dilution buffer. Antibody-bead complexes were washed twice for 5 min rotating at 4°C in 1 ml of each of the following buffers: low salt [50 mM Hepes (pH 7.9), 0.1% SDS, 1% Triton X-100, 0.1% deoxycholate, 1 mM EDTA (pH 8.0), 140 mM NaCl], high salt [50 mM Hepes (pH 7.9), 0.1% SDS, 1% Triton X-100, 0.1% deoxycholate, 1 mM EDTA (pH 8.0), 500 mM NaCl], LiCl [20 mM tris-HCl (pH 8.0), 0.5% NP-40, 0.5% deoxycholate, 1 mM EDTA (pH 8.0), 250 mM LiCl], and TE [10 mM tris-HCl (pH 8.0), 1 mM EDTA (pH 8.0)] using a magnetic rack. Beads were incubated with 250 μl of elution buffer [50 mM tris-HCl (pH 8.0), 1 mM EDTA (pH 8.0), 1% SDS] plus 200 μl of TE with 0.67% SDS for 10 min at 65°C, shaking. RNase A (10 μg) was added and incubated at 37°C for 30 min. Crosslinks were reversed overnight with 40 μg of proteinase K at 65°C. DNA was purified with phenol-chloroform extraction with phase lock tubes followed by ethanol precipitation. DNA was resuspended in 50 μl of 1xTE buffer. ChIP-qPCR reactions were run on a BioRad CFX96 thermocycler with iTaq UniverSYBR Green SMX (BioRad 1725125) in 20 µl reactions and quantitated relative to input dilutions. Starting with genes identified in the rMATs analysis, genes targeted for qPCR had a PSI value of at least 25% and were were enriched for both H3K9me2 and H3K36me3 in previously published datasets (45, 57).

Primers used for ChIP-qPCR analysis:

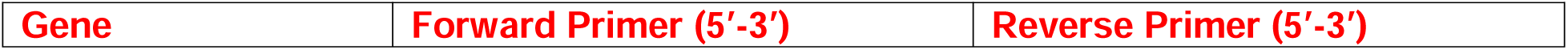
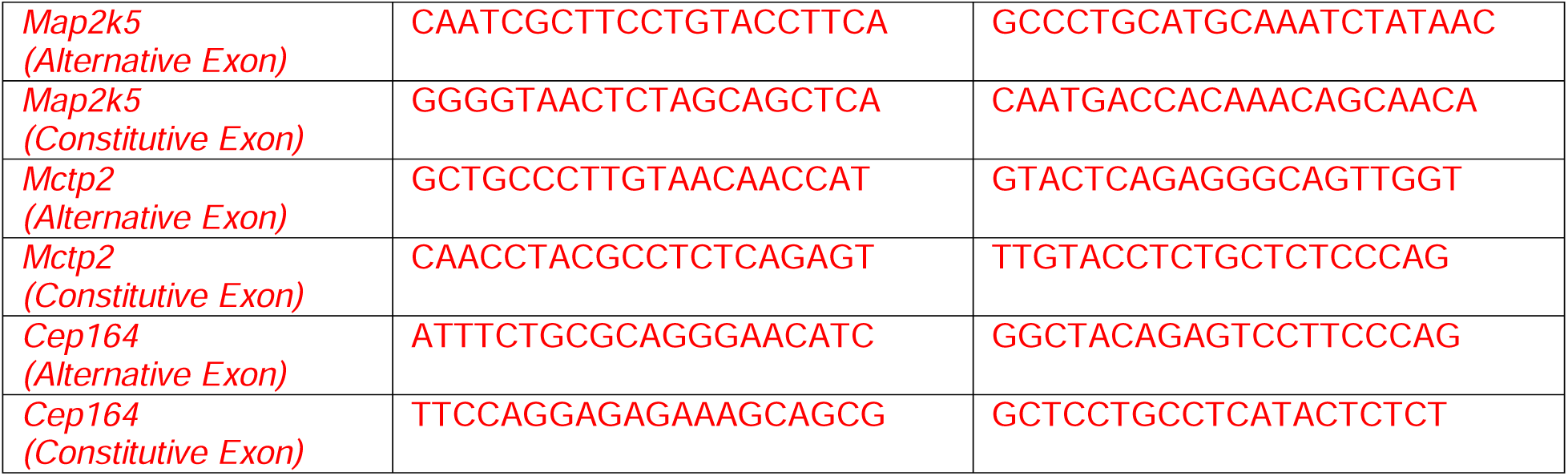

### ChIP-Seq analysis

ChIP-Seq analysis was performed as in as described (56). Briefly, H3K9me2 ChIP-Seq data for WT ESCs with the accession number GSE153651 (58) and GSE177058 (57) were downloaded from Gene Expression Omnibus. Both datasets were in mouse ESCs grown under serum/LIF conditions similar to ours. H3K9me2 ChIP-Seq data for WT and KDM3A/KDM3B DKO ESCs with the accession number DRA006496 was downloaded from the DDBJ database (22). While a different mouse ESC background than we used, this was the only inducible KDM3A/KDM3B double knockout (KO) dataset available to our knowledge. All data was aligned to mm9 using Bowtie2 (59) with the default parameters. Sam files were converted into Bam files and sorted with samtools-1.2 (60) with the default parameters. Peaks were called relative to the input with MACS2 (61) using the following parameters: --broad -p 0.0001. Peak files were annotated using Homer (annotatePeaks.pl). Annotated peaks from both (58) and (57) were overlapped with Venny2.1 (https://bioinfogp.cnb.csic.es/tools/venny/) and used to determine H3K9me2 bound genes in Figure 3. ChIP enrichment was visualized using Integrative Genomics Viewer (IGV) (62). H3K9me2 quantification was calculated from all data sets with bedtools multiBamCov and normalized by read counts per million and per kilobase of gene length.

## RESULTS

### KDM3A and KDM3B interact with splicing machinery in mouse embryonic stem cells

KDM3A and KDM3B have identical catalytic activity and partially redundant functions in several model systems. In mouse embryonic stem cells (ESCs) the deletion of both *Kdm3a* and *Kdm3b* decreases cell number within 4 days (22). KDM3A and KDM3B proteins are conserved in the catalytic Jumonji domain and a zinc finger domain that is required for catalytic activity but have little homology outside of these domains. To gain insight into KDM3A and KDM3B function we performed protein interactome analysis in ESCs. We first generated ESC lines with a FLAG-epitope tagged KDM3A or KDM3B under a doxycycline inducible promoter (Figure 1A and B) and achieved ∼1.5-2 fold overexpression after induction (Figure 1A and B). We then performed immunoprecipitation for each protein from nuclear extracts isolated with benzonase digestion followed by mass spectrometry (Figure 1B, Supplementary Figure S1A). After stringent filtering for common contaminants of FLAG pulldowns (Crapome database (43) Saint score 0.7, Methods), we identified 122 proteins that were commonly enriched over background uninduced controls in both the KDM3A and KDM3B interactomes and 37 and 35 unique interactions respectively (Figure 1C). We further classified the interactions using STRING network analysis (Figure 1D, Supplementary Figure S1B and S1C). KDM3A and KDM3B demethylate the repressive H3K9me2 that is enriched in heterochromatin that would ultimately result in an increase in transcription. Therefore, we expected to find interactors that are transcriptional activators. Instead, we found associations with repressor proteins including the Heterochromatin Protein 1 family (CBX1,CBX3, CBX5), DNA methyltransferase complex - DNMT3A and DNMT3L, components of the polycomb complex-EED, histone deacetylase HDAC6 and components of the nuclear lamina such as BANF1 (Figure 1D).These associations suggest that KDM3A and KDM3B are localized to a heterochromatin environment within the cell where they may initiate the first step of derepression. Surprisingly, the majority of KDM3 interacting proteins belonged to the RNA processing machinery (Figure 1C and D) along with the FACT and PAF complexes that associate with actively elongating RNAPII. The RNA processing proteins included members of the spliceosomal machinery, which removes introns from pre-mRNA (63), and RNA helicases, which are important for general RNA metabolism and rearrangement of ribonucleoprotein complexes (64). We also found components of the exon junction complex, which control post-transcriptional export and stability of mRNAs (65) (Figure 1D and E).

**Figure 1:**
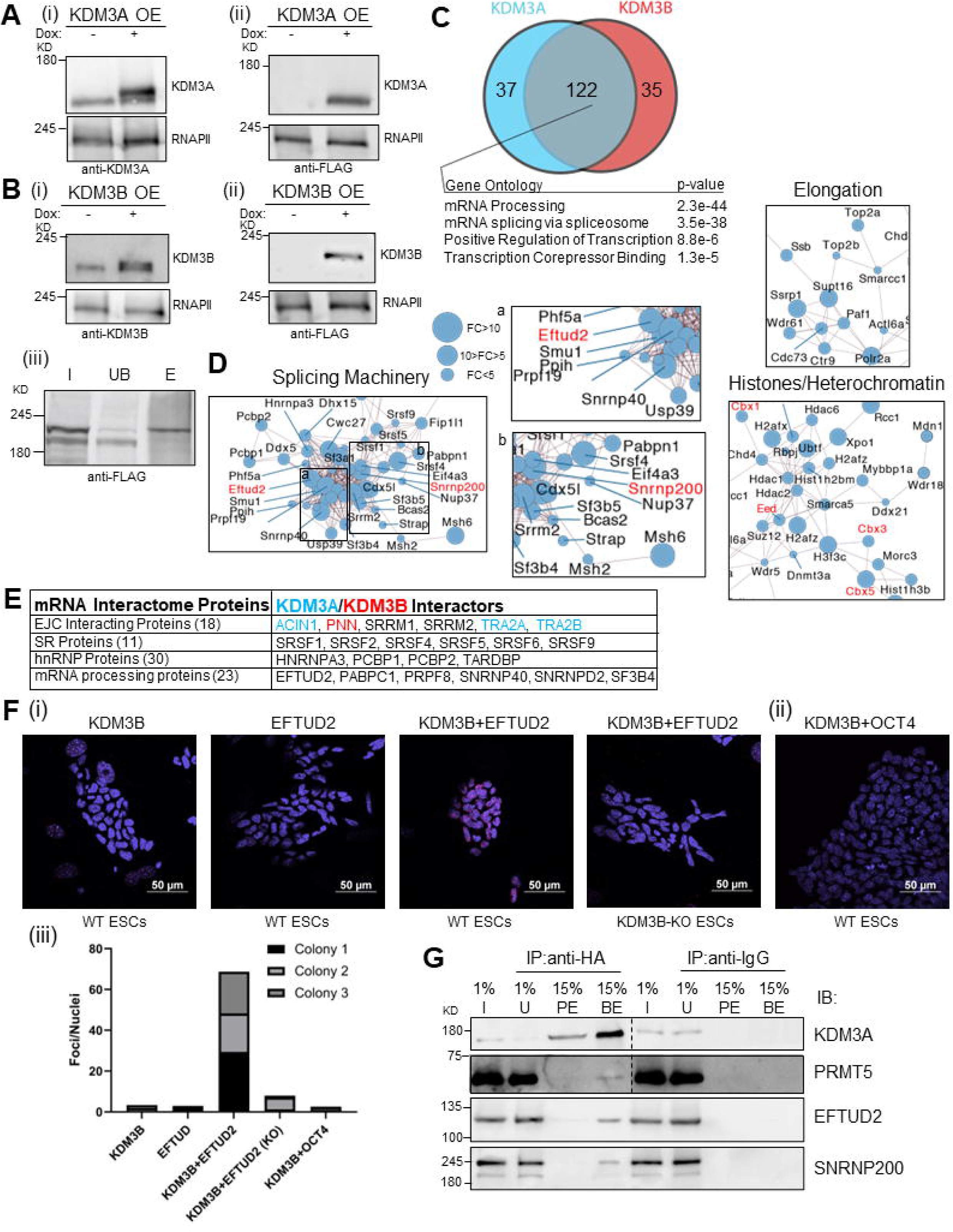
KDM3A and KDM3B interact with splicing machinery in ESCs. (A) Immunoblot of (i) KDM3A, (ii) FLAG in ESCs with doxycycline (dox)-inducible expression of FLAG-KDM3A. Loading control = RNA polymerase II (RNAPII). OE=Overexpression (B) Same as above except in dox-inducible FLAG-KDM3B ESCs (i) KDM3B (ii) FLAG (iii) FLAG following immunoprecipitation with FLAG antibody. I=Input, UB=Unbound, E=Elute (C) Venn diagram of KDM3A and KDM3B interactome. Filtered for nuclear compartment using DAVID GO (https://david.ncifcrf.gov/) analysis and frequent FLAG pulldown contaminants using “Crapome” (https://reprint-apms.org/) repository at SAINT score of 0.7 (43). Gene Ontology (GO) enrichment of shared interactome are listed at enrichment score>2.5 and p-value<.00001. (D) String network analysis (https://string-db.org/) of KDM3A and KDM3B shared interactome with high confidence interaction scores (0.7 minimum) from database and experimental sources. Node size represents fold change (FC) of KDM3A over uninduced spectral counts. FC>10 = large node, 10>FC>5 = medium node, FC<5 = small node. Inset a and b included for better view of interacting splicing machinery. A few small node protein labels are removed to improve legibility. See Supplementary Table 1 for all interacting proteins. (E) KDM3A and KDM3B interactors overlapped with functional categories of mRNP components from Singh et al. (119) or categorized as other mRNA processing proteins. Shared= black, KDM3A unique=blue, and KDM3B unique=red. (F) Single slice of 1 uM confocal images of *in situ* Duolink™ Proximity ligation assay (PLA) of (i) KDM3B single antibody, EFTUD2 single antibody, and KDM3B+EFTUD2 antibodies in WT ESCs or KDM3B+EFTUD2 antibodies in KDM3B KO ESCs. (ii) PLA of KDM3B and OCT4 in WT ESCs. Scale bar = 50µM with 60x objective. (iii) PLA foci and nuclei were quantified in ImageJ of one colony per field of view. The ratio of foci per nuclei is reported for each colony. (G) Immunoprecipitation (IP) with HA antibody or mouse IgG with immunoblot (IB) for KDM3A, SNRNP200, PRMT5, and EFTUD2. I=Input, U=Unbound, PE=Peptide Elute, BE=Boil Elute. Dotted line represents a cropped image.

To validate these interactions in a system that did not overexpress either protein, we inserted a HA epitope tag into the endogenous locus of both KDM3A and KDM3B. From co-immunoprecipitation experiments, we confirmed an interaction of KDM3A with PRMT5, an arginine methyltransferase that has spliceosomal substrates and effects on splicing in ESCs (66). We also validated an interaction (albeit at substoichiometric levels) with two protein components of the U5 snRNP: the GTPase EFTUD2, and SNRNP200, which is involved in spliceosome assembly, activation and disassembly (67) (Figure 1G and Supplementary Figure 1D.

Spliceosomal proteins are far more abundant than KDM3A or KDM3B (Supplementary Figure S1D), hence to further validate these interactions under physiological conditions in situ, we performed a proximity ligation assay (PLA) (Methods, Duolink™). This assay detects protein interactions that are less 40 nm apart. We found a highly specific interaction signal, represented by red foci, only in the presence of both KDM3B and EFTUD2 antibodies. This interaction was absent when cells were stained with either antibody alone (Figure 1F), and in an ESC line that was knocked out for KDM3B (Figure 1F, Supplementary Figure S1E). Further, the abundant ESC specific nuclear protein OCT4 (POU5F1), that was not detected in the KDM3A/KDM3B interactome, did not show an interaction by proximity ligation assay (Figure 1F). KDM3B has been reported to associate more robustly with transcriptionally active chromatin (58). We inhibited cells with 5,6-Dichloro-1-beta-Ribo-furanosyl Benzimidazole (DRB) that pauses RNA polymerase II and repeated the PLA. We found the relative association of KDM3B with EFTUD2 decreased upon DRB treatment suggesting that KDM3B splicing associations can be modulated but are not dependent on ongoing transcription (Supplementary Figure S1F). Taken together we uncover an interaction of the histone demethylase KDM3A and KDM3B proteins with the mRNA processing machinery.

### Delayed accumulation of H3K9me2 upon acute degradation of KDM3A and KDM3B proteins

The majority (∼80% in human cells) of mRNA processing is thought to occur co-transcriptionally, with some variation between species and cell types (68–71). Most active spliceosomes are associated with chromatin (72). Traditional methods of genomic deletion such as conditional genomic deletion or transcriptional intervention with RNAi knockdown would only remove the KDM3A and KDM3B proteins slowly, confounding direct and indirect effects on splicing. Therefore, we needed a model of rapid and inducible protein degradation. We gene-edited an inducible “degron” tag into the endogenous locus of both KDM3A and KDM3B (Figure 2A). To facilitate CRISPR-Cas9 mediated gene editing we used microhomology mediated end joining (MMEJ) (32) (Supplementary Figure S2A) to insert the degron tag into the 5’ end of each KDM3 locus to avoid the 3’ end that encodes the Jumonji catalytic domain. The degron tag contains a 2xhemagglutinin (HA) tag and is derived from a mutated variant of the FKBP12 protein, FKBP12^F36V^, which responds to the dTAG-13 ligand for targeting the tagged protein to the ubiquitin proteasome pathway (31). We initially obtained clones that were degron tagged at both *Kdm3b* alleles with decreased expression of the KDM3B protein (Figure 2B). Even though depletion of KDM3B has been shown in some cases to lead to genome instability (73), importantly, these clones maintained a stable karyotype throughout the derivation process (Supplementary Figure S2C). The Kdm3b-degron tagged clones were then targeted with the *Kdm3a* degron construct and heterozygous clones were obtained where one allele of *Kdm3a* codes for an in-frame degron tag and the second allele was knocked-out by an insertion at the 3’ repair site (Supplementary Figure S2B and D). Both KDM3A and KDM3B proteins were completely degraded upon exposure to the small molecule dTAG-13 (Figure 2B). From a timecourse of dTAG-13 exposure we determined that KDM3A/KDM3B could be degraded as soon as 15 minutes (Figure 2C) and was completely undetectable even at 24 hours after a single application of the dTAG-13 molecule (Supplementary Figure S2E). By contrast, the target of KDM3A/3B catalytic activity, H3K9me2, is relatively stable at the global level over 48 hours of degradation, with minimal gain at 24 hours compared to the uninduced condition (Figure 2D). H3K9me2 could be quickly processed to H3K9me3. However, H3K9me3 also remained relatively stable over the 48-hour timecourse (Figure 2D).

**Figure 2:**
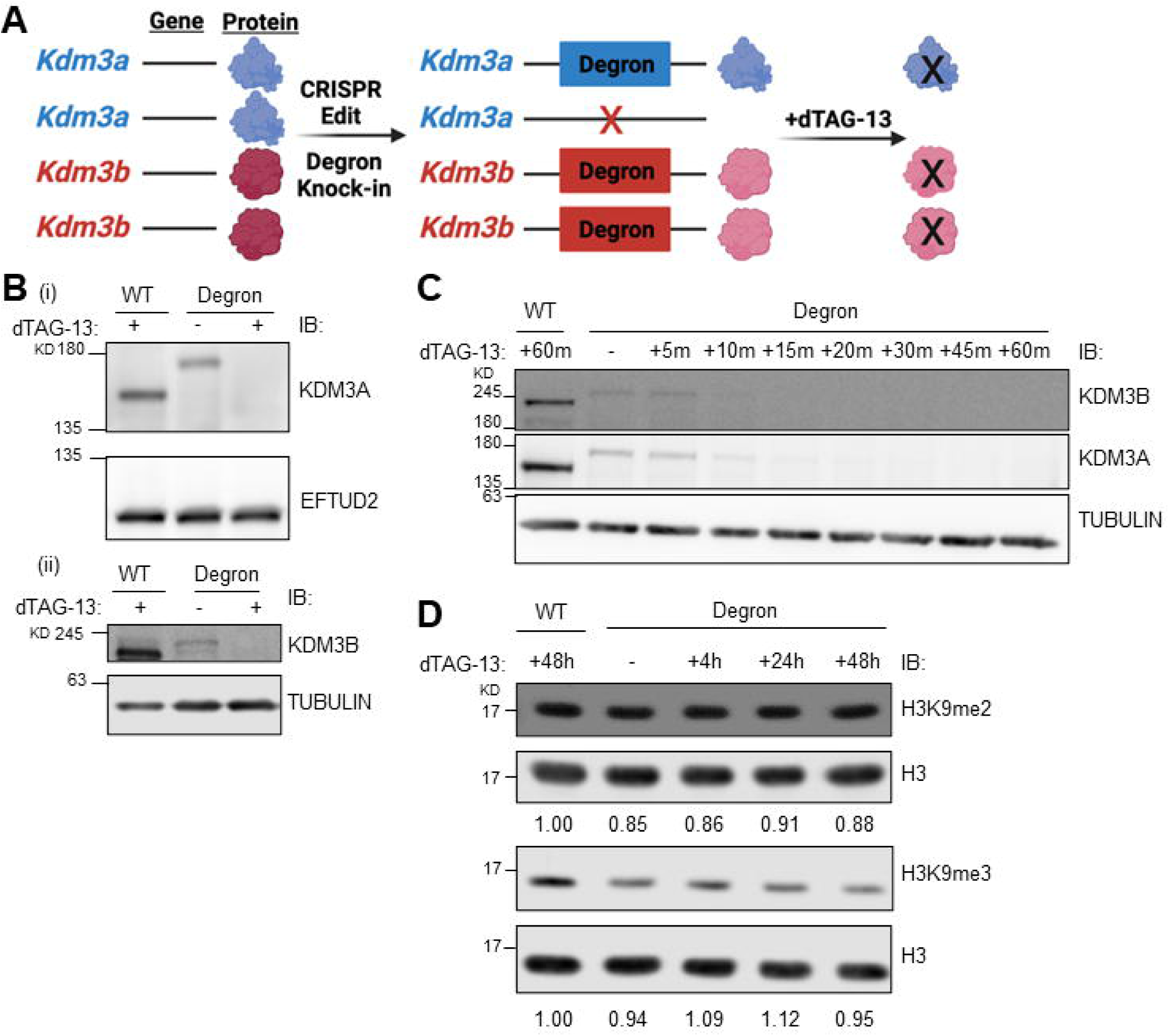
Inducible double knockout of KDM3A and KDM3B results in delayed global accumulation of H3K9me2. (A) Schematic of gene editing strategy to generate degron inducible KDM3A and KDM3B proteins. (B) Immunoblot for (i) KDM3A or (ii) KDM3B in WT ESCs or KDM3A/KDM3B degron ESCs. Note increased size of degron tagged protein. dTAG-13 ligand induced samples (+) treated for 24 hours. (i) Loading control= EFTUD and (ii) Loading control = TUBULIN (C) Immunoblot for KDM3A, KDM3B, and TUBULIN (loading control) in less than one hour of dTAG-13 induced degradation. (D) Immunoblot of H3K9me2, H3K9me3, and H3 (loading control) in WT ESCs and degron ESCs at indicated timepoints of degradation.

We performed immunofluorescence (Supplementary Figure S2F) and proximity ligation assays with HA (targeting both KDM3A and KDM3B) and EFTUD2. We detected PLA foci between the proteins of interest and observed a decrease upon dTAG-13 mediated degradation of KDM3A and KDM3B (Supplementary Figure S2G). Thus, using the degron strategy established a system in which we could study the direct effects of KDM3A and KDM3B loss in a timescale appropriate for splicing defects.

### Complete degradation of KDM3A and KDM3B leads to differential gene expression agnostic of H3K9me2 status

We performed RNA-seq of the degron clones at 4 hours after exposure to dTAG-13 to capture immediate changes in both gene expression and splicing. The target of KDM3A and KDM3B histone demethylase activity, H3K9me2, is enriched in broad megabase domains that repress several development specific genes (17). However, upon complete loss of all KDM3A/KDM3B protein we found only 420 genes with differential expression (DEG) compared to wildtype ESCs (Figure 3A) (Supplemental Table 2). The proteins in the shared KDM3A and KDM3B interactome are not direct transcriptional targets since they were not differentially expressed after KDM3A/KDM3B degradation (data not shown). The upregulated and downregulated genes were enriched for functional categories of cell migration and signaling pathways; and interferon response respectively (Figure 3A). Despite this functional distinction both the up and down-regulated DEG were lower expressed as compared to the global median of genes in ESCs (Figure 3B).

**Figure 3:**
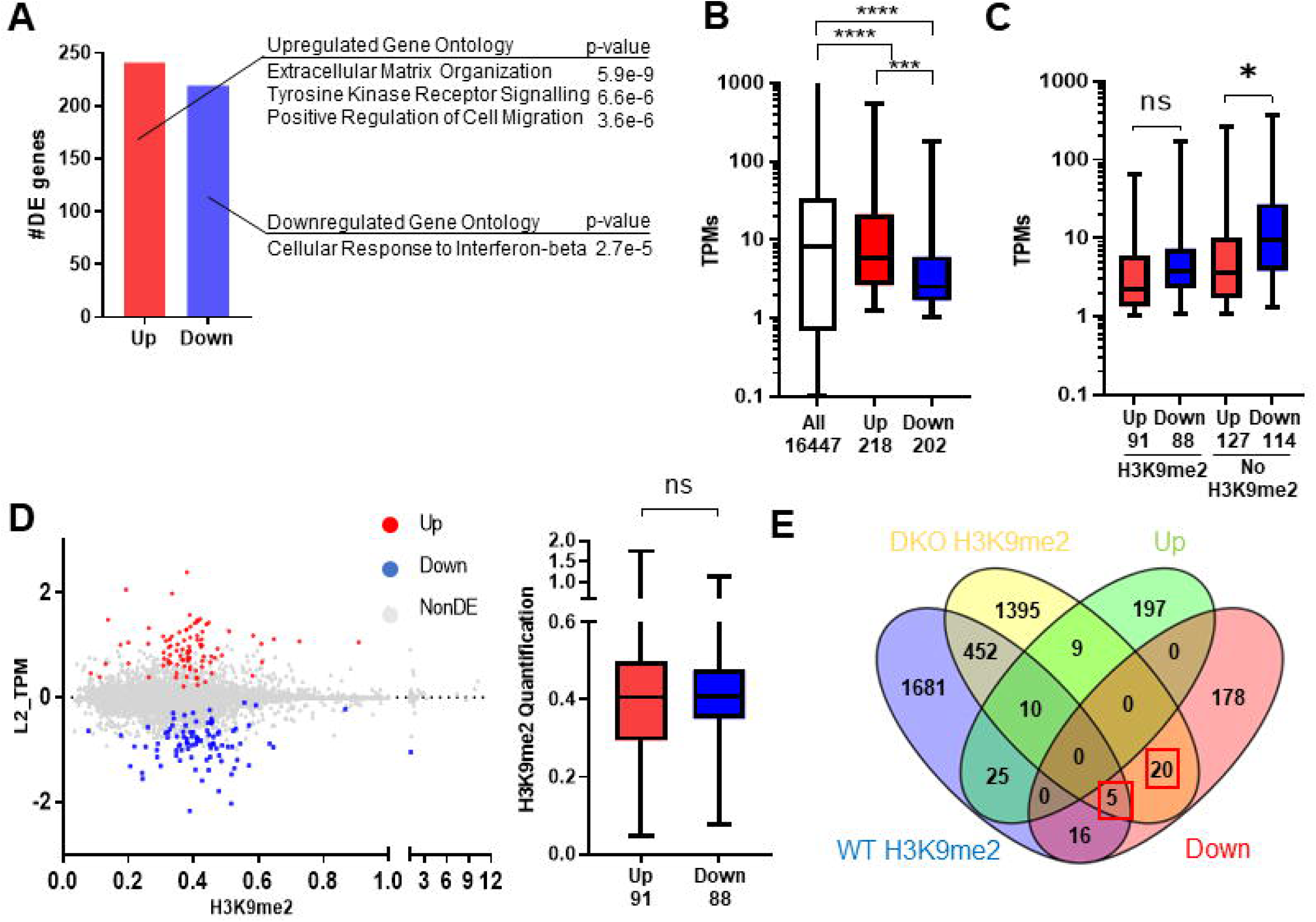
Acute degradation of KDM3A and KDM3B leads to differential gene expression agnostic of H3K9me2 status. (A) Number of differentially expressed genes (DEG) between Double Degron (DDeg+) exposed to dTAG-13 ESCs and wildtype ESCs. Gene ontology categories for DEGs shown at minimum p-value < 0.00001. (B) Boxplot of absolute expression (TPMs) in ESCs for all annotated genes from RSEM quantification with a non-zero median TPM (All expressed genes), TPMs from up or down DEGs. Numbers indicate number of genes per condition. ****p-value < 0.0001 assessed by unpaired Welch’s t-test. (C) Boxplot of average TPMs+1 from DEGs with (H3K9me2) or without (No H3K9me2) a called consensus H3K9me2 peak (57, 58). *p-value < 0.05 assessed by unpaired Welch’s t-test (D) Left: Scatterplot of log2 (DDeg+/WT) fold change of TPMs of Non-DE and DEG against H3K9me2 enrichment (normalized read counts per million and by 1kb per gene) at those genes. Right: Boxplot of total H3K9me2 enrichment at up or down DEG. (E) Four-way Venn overlap of DEG and H3K9me2 peaks in WT ESCs and KDM3A and KDM3B DKO ESCs (22).

To determine if the low expression of DEGs was because of prevailing H3K9me2, we overlapped the DEGs with consensus peaks from two published H3K9me2 chromatin immunoprecipitation sequencing (ChIP-seq) data sets (57, 58). A significant proportion of the DEGs (179 genes) were enriched for H3K9me2, indicating that H3K9me2 may participate in localizing KDM3A and KDM3B to affected genes (Supplementary Figure S3A). Irrespective of whether they were up or downregulated upon KDM3A/KDM3B degradation, the DEGs that had a H3K9me2 peak were lower expressed than those that lacked H3K9me2 (Figure 3C). Among the genes that contained a H3K9me2 peak, the overall H3K9me2 enrichment was not significantly different between up and down DEG (Figure 3D).

Finally, to determine the genes that may be directly regulated by the loss of KDM3A and KDM3B, we analyzed a third published set of H3K9me2 ChIP-seq containing KDM3A/KDM3B DKO ESCs and WT ESCs (22). Assuming loss of the two demethylases would lead to the spread of repressive H3K9me2, we reasoned genes that were downregulated and bound by H3K9me2 in the DKO ESCs could be direct targets of the enzymes and found 25 genes that fit this criterion (Figure 3E). We did not find a significant difference between the H3K9me2 quantification of DEG genes in the WT or DKO ESCs (Supplementary Figure S3B). Taken together with the lack of change in global H3K9me2 levels (Figure 2C) and the small number of DEG that are enriched for H3K9me2 at 4 hours (Figure 3E), we conclude that these gene expression changes are not dependent on H3K9me2 status but may instead correlate with the presence of KDM3A or KDM3B proteins. Upon the degradation of KDM3A or KDM3B, essential protein complexes that are scaffolded or nucleated by KDM3A or KDM3B may be disrupted, leading to changes in gene expression or alternative splicing.

### Alternative splicing defects upon acute loss of KDM3A/KDM3B occur in signaling and chromatin related genes

Due to the interaction of KDM3A and KDM3B with splicing machinery, we analyzed the RNA sequencing for splicing aberrations using rMATs. rMATS detects differential splicing events by modeling exon inclusion versus exon skipping levels in multiple replicates to calculate a ‘percent spliced in (PSI)’ for each exon considered (52, 74). rMATs reports canonical splicing events such as skipped exons, alternative 5’ and 3’ splice site selection, and mutually exclusive exons (Figure 4B). We performed rMATS with cutoffs of FDR < 0.1, PSI difference of 5%, and a 15 read minimum for each junction and identified 901 significant events within 729 genes (Figure 4B).

**Figure 4:**
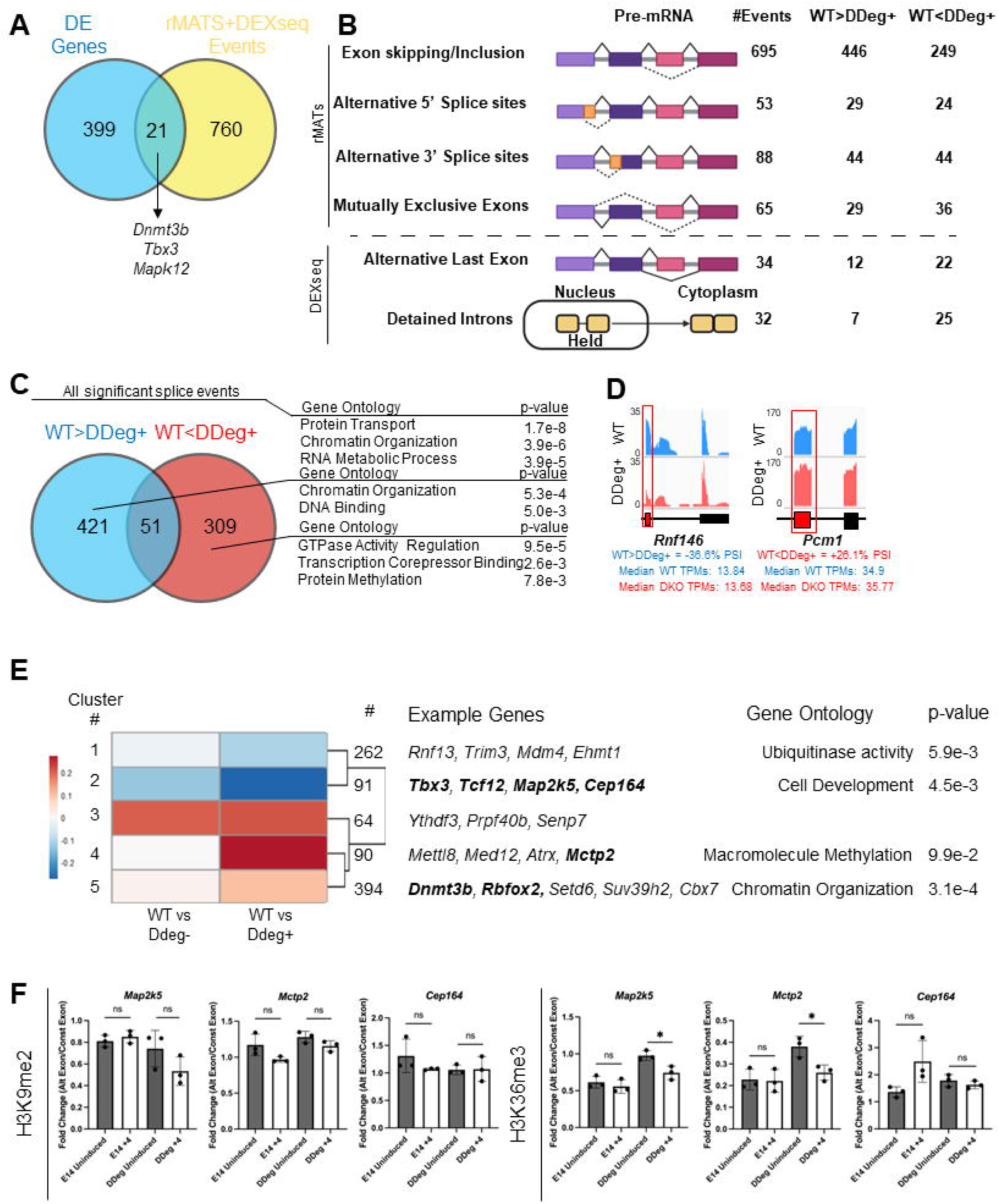
Alternative splicing defects upon acute loss of KDM3A/KDM3B occur in signaling and chromatin related genes. (A) Venn overlap of DE Genes (blue) and aberrantly spliced genes (yellow) called from rMATs and Dexseq analysis. (B) Left panel: schematic of alternative splicing pathways, results above the dotted line are from rMATs and below are from Dexseq. Right panel: Number of significant events called in each class of alternative splicing pathway, FDR<0.1, inclusion level difference x>+/-0.5, and 15 read minimum. (C) Venn overlap of aberrantly spliced genes sorted by higher inclusion in WT (blue) or in DDeg+ (red). Gene ontology categories for aberrantly spliced genes shown, minimum p-value < 0.001. (D) IGV tracks for example genes of exons with higher percent spliced in (PSI) for WT or DDeg+ conditions. PSI value calculated in rMATs from 3 replicates in each condition. Negative value indicates event is included more in the WT condition and vice versa. The affected exon is marked in red. (E) Clustering (k-means=5) of differentially spliced events by inclusion level difference score generated in rMATs. WT vs uninduced degron (DDeg-) and WT vs induced degron (DDeg+) lines are shown. A positive number indicates greater inclusion in WT cells. Example genes from clusters and significant gene ontology terms (minimum p-value < 0.001) are in right panel. (F) ChIP-qPCR of WT (E14) and inducible KDM3A/KDM3B double degron (DDeg) ESCs uninduced or treated for 4 hours with dTAG-13. Left, H3K9me2 ChIP-qPCR. Right, H3K36me3 ChIP-qPCR. Three technical replicates performed for each condition which include alternate and constitutive spliced exons. Fold change determined by dividing alternative exon replicate over constitutive exon replicate.

KDM3A/KDM3B interact with PRMT5, a known regulator of non-canonical splicing called detained introns in several cell types (75). Detained introns refer to mRNA containing one or more introns that are trapped in the nucleus and ultimately degraded, or post-transcriptionally spliced and exported into the cytoplasm in response to additional cues. While PRMT5 was strongly enriched in the KDM3A interactome (Supplementary Table 1), it is also a common contaminant in FLAG-mediated immunoprecipitation mass spectrometry (Crapome (43).

Therefore, we first validated the interaction with PRMT5 independently using an HA-epitope that is introduced along with the degron domain in the KDM3A/KDM3B degron ESC line. We found a HA-KDM3A/KDM3B interaction with PRMT5 using PLA that was lost upon the addition of dTAG-13. (Supplementary Figure S4A). An endogenous KDM3B-PRMT5 interaction was also observed in a second, feeder-dependent ESC line (Supplementary Figure S4B).

We performed DEXseq (55) analysis to identify DIs. DEXseq calculates the ratio between the read counts of a particular exon and the counts at the exons at the rest of the gene to calculate relative exon usage (55). By combining DEXseq with a custom annotation including detained introns (54) we assessed the splicing effects upon the acute loss of KDM3A and KDM3B. At a DEXseq threshold of padj < 0.1 we identified several genes that had detained introns (32 events), as well as some with alternative polyadenylation sites (34 events) (Figure 4B).

Combining the significant events from the rMATS and DEXseq analyses, we found the splicing of 781 genes to be impacted by loss of KDM3A/KDM3B (Supplemental Table 3) but only 21 genes were both differentially expressed and changed in splicing patterns (Figure 4A) suggesting that the changes in isoforms do not immediately lead to expression changes (76). We found a greater number of exon skipping/inclusion events (446) with a negative PSI (WT>DDeg+) than positive PSI (WT<DDeg+, 249), insinuating the loss of KDM3A and KDM3B leads to more exon skipping. Other than a 3-fold increase in detained introns in the DDeg+ condition, we found the directionality within other event classes to be evenly distributed between the WT and DDeg+ conditions (Figure 4B).

We then evaluated whether the genes affected in the DDeg+ condition, were enriched for specific functions through DAVID GO analysis. While all KDM3A/3B responsive genes, regardless of PSI direction, had a significant enrichment for gene ontology terms such as Protein Transport, Chromatin Organization, and RNA metabolic Processing (Figure 4C), genes with a positive PSI value (309 genes in WT<DDeg+) were associated with GTPase Activity Regulation, Transcription Corepressor Binding, Protein Methylation (Figure 4C, 4D). By contrast, genes with a negative PSI value (421 genes in WT>DDeg+) were associated with Chromatin Organization and DNA Binding (Figure 4C, 4D). The largest class of splicing event, skipped exons (SEs), included several proteins that had sequence-specific DNA binding such as transcription factors *Tbx3, Meis1,* and *Zfp263.* To gain further insight into the splicing events that were most susceptible to the acute loss of the KDM3 proteins, we analyzed the 901 significant canonical splicing events found in the WT/DDeg+ analysis and compared this to the PSI values between WT and the DDeg ESCs prior to the addition of the dTAG-13 ligand (WT vs DDeg-), which have slightly reduced expression of KDM3A and KDM3B and performed k-means clustering into 5 clusters (Figure 4E).

In general, most of the splicing events were further exacerbated upon complete degradation of KDM3A and KDM3B. Cluster 3 was agnostic to the dTAG-13 addition included genes that function in RNA processing such as *Ythdf3* and *Prp40b*. Chromatin-associated proteins such as *Atrx* and *Med12* and chromatin-modifying enzymes *Dnmt3b*, *Setd6* and *Suv39h2* had splice events included more in the WT cells (WT>DDeg+) (Figure 4E). *Rbfox2,* an RNA-binding protein that regulates alternative splicing events, was also in the WT>DDeg+ group (Supplementary Figure S4C). The genes with splice events that were more included in DDeg+ cells (WT<DDeg+) functioned in ubiquitinase activity such as *Rnf13, Rnf214 and Trim3*. Cluster 2 was most sensitive to the amount of KDM3 proteins included kinases in the MAP pathway *Map2k5* and transcription factors such as *Tbx3* and *Tcf12* that are important for pluripotency.

Although globally there was minimal difference in the abundance of H3K9me2, we wondered about the status of this modification at the alternatively spliced exons. We performed chromatin immunoprecipitation for H3K9me2 and as a control for H3K36me3, which is implicated in splicing (77–80) on WT ESCs (E14) and KDM3A/KDM3B degron (DDeg) ESCs treated with dTAG-13 for 4 hours. We amplified genes that were enriched for both H3K9me2 and H3K36me3 in previously published datasets (45, 57). The affected exon from rMATs (alternative exon) and a neighboring exon not detected in rMATs and not sharing a junction with the affected exon (constitutive exon) were targeted for qPCR at *Map2k5, Mctp2,* and *Cep164* (Supplementary Figure S4D). Across two biological replicates, we found small, mostly nonsignificant and inconsistent changes in H3K9me2 and H3K36me3 at these exons after four hours of degradation in the presence of dTAG-13, indicating that there is not an obvious chromatin effect that is driving the splicing changes upon KDM3A and KDM3B degradation (Figure 4F and Supplementary Figure S4E). Considering that these genes were marked with H3K9me2, however, the chromatin modifications could be serving as adaptors to assist in the targeting of splicing machinery by the KDM3 enzymes.

### Rescue of alternative splicing defects caused by loss of KDM3A/KDM3B does not depend on catalytic activity of KDM3A

We next sought to determine if the splicing changes seen upon KDM3A and KDM3B degradation were dependent on the catalytic activity of these enzymes. To answer this question, wildtype and two independent DDeg ESCs were transduced with a lentivirus expressing either full length FLAG-wildtype (FL) or FLAG-catalytic mutant (CM) of KDM3A (35) and selected for 7-10 days (Figure 5A). WT-KDM3A or CM-KDM3A was robustly expressed in both the wild type ESCs and DDeg ESCs with rescue clone #1 having slightly less FL expression and rescue clone #2 slightly less CM (Figure 5B). After treatment of each ESC population with dTAG-13 for 4 hours, there was a decrease in the endogenous KDM3A in the DDeg ESCs but as expected not in the FLAG-WT-KDM3A or FLAG-CM-KDM3A.

**Figure 5:**
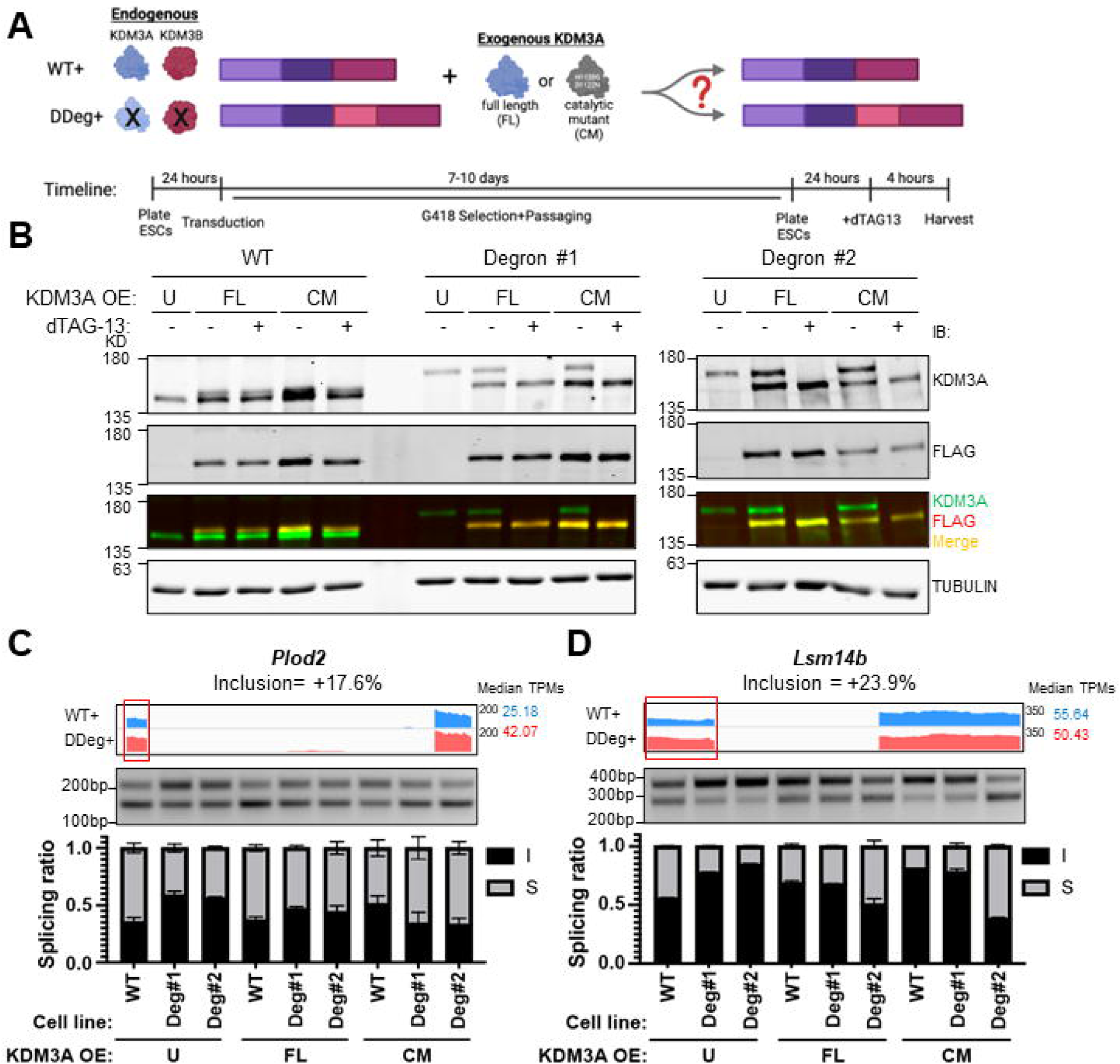
Rescue of alternative splicing defects caused by loss of KDM3A/KDM3B does not depend on catalytic activity of KDM3A. (A) Schematic (top) and timeline (bottom) of KDM3A full length (FL) or KDM3A catalytic mutant (CM) overexpression rescue experiment. (B) Immunoblot for KDM3A, FLAG, and TUBLUIN (loading control) in WT ESCs or Degron clones untransduced (U), transduced with full length KDM3A (FL), or transduced with catalytic mutant KDM3A (CM). (C) Semi-quantitative PCR of splicing targets in WT ESCs and Degron clones at 4 hours of degradation, each untransduced (U), transduced with full length KDM3A (FL), or transduced with catalytic mutant KDM3A (CM). Representative IGV tracks depicting affected exon (red box) are below. (D) Electrophoresis image from 5C was quantified using Licor software, fluorescence intensity reported as arbitrary units (a.u.) and inclusion/skipped ratio calculated and represented in right panel. I=included ratio and S=Skipped ratio.

At selected target genes from the rMATs analysis (Methods) we found the four-hour degradation of KDM3A and KDM3B led to the increased inclusion of an exon in both *Plod2* and *Lsm14b* in both the DDeg ESC clones as compared to the WT ESCs (Figure 5C and 5D). When compared to untransduced WT ESCs, the overexpression of either FL-KDM3A or CM KDM3A rescued expression of the affected exons by reducing inclusion to levels on par with the wildtype condition irrespective of slight differences in FL and CM expression between clones (Figure 5B-5D). This indicates that the catalytic activity of these enzymes is not necessary for their role in regulating splicing.

### KDM3A/KDM3B depletion leads to the missplicing of regulators of the naïve state of pluripotency

Mouse pluripotent states exist in a continuum of pluripotency ranging from the “ground state” that resembles the early blastocyst to a “naïve state” resembling early epiblast and “primed state” obtained from the late epiblast. The ground state ESCs can be obtained from embryos by deriving ESCs in 2i/LIF conditions without serum while the “naïve state” can be captured in serum/LIF conditions. ESCs can be converted from the naïve state (in which all our experiments were performed) to ground state by passaging in 2i/LIF conditions (Figure 6A). We were struck by the differential splicing of several regulators of the pluripotency continuum in our dataset. To investigate this idea further we evaluated splicing in the ground state of pluripotency.

**Figure 6:**
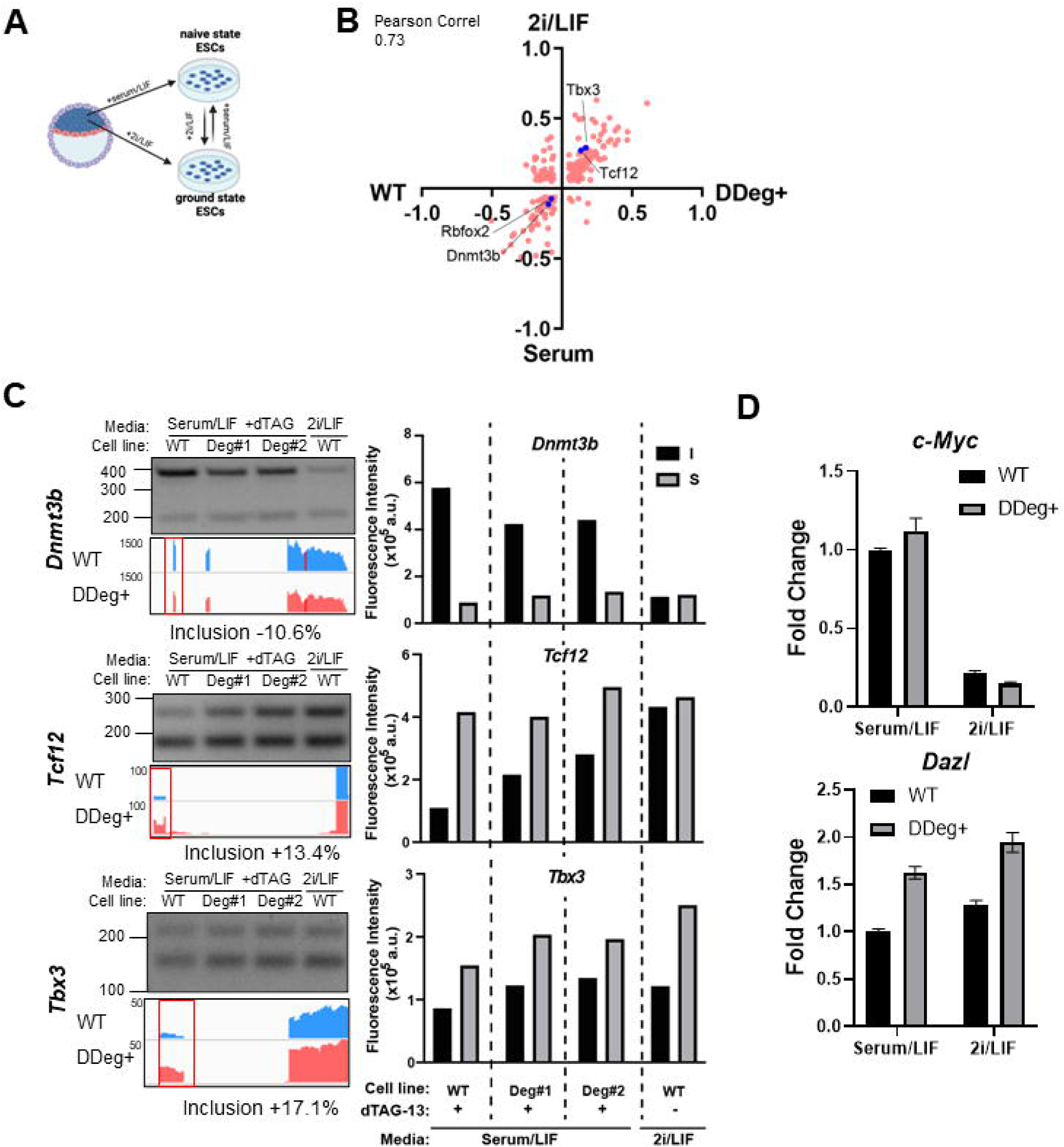
Loss of KDM3A and KDM3B leads to differences in stable naïve state. (A) Schematic detailing derivation of naïve ESCs (serum/LIF) or ground ESCs (2i/LIF). (B) Scatterplot of splicing events called by rMATs in both WT vs DDeg+ ESCs as well as serum vs 2i/LIF ESCs (82) measured by inclusion level difference score. Pearson correlation between the inclusion level difference of specific events called in both comparisons. Blue dots represent events chosen for further confirmation. (C) Semi-quantitative PCR of splicing targets in WT ESCs, DDeg+ clones at 4 hours of degradation, and 2i/LIF ESCs. Representative IGV tracks depicting affected exon (red box) are below. Electrophoresis image was quantified using Licor software, fluorescence intensity reported as arbitrary units (a.u.) and represented in right panel. I=included isoform and S=Skipped isoform. (D) Relative expression of c-Myc or Dazl in WT or DDeg+ ESCs in serum/LIF or 2i/LIF conditions. WT cells in serum/LIF set to 1.

We performed rMATS analysis to compare canonical splicing events between our WT ESCs which were cultured in serum/LIF and ESCs grown in 2i/LIF conditions from two sets of published data (Methods). There were ∼4000-5000 splicing differences detected between serum/LIF and 2i/LIF conditions indicating that splicing patterns change globally between these cell fates (Supplementary Figure S5A and B). We then asked how similar the direction of splicing differences upon KDM3 degradation were to ESCs grown in 2i/LIF conditions. There was a greater overlap between splicing events that were gained in the Double degron or 2i/LIF conditions compared to WT ESCs grown in serum/LIF (Supplementary Figure S5A and B).

When comparing the magnitude of changes, we found a large positive correlation in the inclusion levels of splice forms between the DDeg+ and the 2i/LIF conditions (Pearson = 0.73 with Kinoshita et al. (81) and 0.51 with Yang et al. (82) (Figure 6B and Supplementary Figure S5C). Therefore, upon degradation of the KDM3 proteins the alternative splicing of genes trends toward a more ground pluripotent state pattern.

Among the splicing events we focused on candidates that have been implicated in cell fate switches in pluripotency and were similarly spliced in the WT-DDeg+ and, WT-2i/LIF comparison. *Dnmt3b*, a de novo methyltransferase that is lowly expressed in ground state cells and gains in expression toward primed pluripotency. The different isoforms of Dnmt3b that are detected in our data lack two exons that code for conserved motifs within the catalytic domain and are thought to be catalytically inactive (83–85). *Tbx3* is a transcription factor with two isoforms that regulate the expression of pluripotency gene *Nanog* (86). *Tcf12* is a transcription factor whose deletion compromises mesodermal specification and reduces the expression of *Nanog* and other transcription factors in ESCs and can destabilize self-renewal capacity (87, 88).

To validate our results, we performed semi-quantitative RT-PCR analysis for three candidate genes in two independent clones of KDM3A/KDM3B with degron tags and in WT ESCs grown in serum/LIF and 2i/LIF conditions (Figure 6C and Supplementary Figure S5D). In every case we found that the splicing pattern in KDM3A/KDM3B degraded clones trended towards the ESCs grown in 2i/LIF conditions, indicating a shift towards ground state.

The transcriptomes of serum and 2i/LIF ESCs are highly similar (89). Among the few genes that are significantly different between the two conditions, we found that genes higher in 2i/LIF, such as *Dazl*, were slightly increased in the double degron cell lines, while genes higher in serum/LIF, such as *c-Myc*, were not downregulated (Figure 6D). Notably, the exposure of double degron cell lines to 2i/LIF conditions increased *Dazl* levels compared to WT cells and decreased c-Myc levels further than in WT cells (Figure 6D). Taken together we conclude that altering KDM3 levels impacts important regulators of cell fate by splicing leading to a compromised naïve state of pluripotency.

## DISCUSSION

In this study, we have found the histone demethylases, KDM3A and KDM3B, to be important regulators of RNA splicing in mouse ESCs (mESCs) independent of the H3K9 methylation status of target genes. Histone demethylases have a profound impact on processes such as development, gene regulation, and disease manifestation (90). The primary function of histone demethylases is in modulating transcriptional activity by altering local chromatin structure and providing a platform for the recruitment of histone modification “reader” proteins that transduce downstream effects. Histone demethylases can also have non-histone targets, expanding their range of action (91, 92). Beyond these catalytic activities, histone demethylases are members of larger protein complexes that modulate their activity, raising the possibility of non-catalytic functions such as scaffolding. In pluripotent stem cells KDM3A and KDM3B are implicated to have overlapping function, since only the deletion of all four alleles induces lethality in mESCs and developing epiblasts (22). Supporting this idea of redundant function we have found that the majority of KDM3A and KDM3B protein interactors are the same. That most of these overlapping binding partners were related to RNA splicing machinery implicates KDM3A and KDM3B function in post-transcriptional control in addition to their traditionally acknowledged transcriptional roles in pluripotency. These effects of KDM3A and KDM3B are independent of their catalytic function and H3K9me2 status, even though prior studies have shown that both H3K9me2 and H3K9me3 across exons affects the kinetics of splicing and can lead to increased exon inclusion by affecting RNA pol II processing (93–96).

Splicing misregulation in ESCs can lead to loss of self-renewal, differentiation defects and cell death (97–99). In human ESCs, only specific spliced isoforms of the master pluripotency transcription factor OCT4 can support self-renewal (100). During somatic cell reprogramming the entire pattern of splicing is switched from that of a somatic cell to a pluripotent cell (101). A specific isoform of FOXP1 generated through alternative splicing has been shown to promote the maintenance of ESC pluripotency and contribute to efficient reprogramming of somatic cells (102). Beyond these examples of transcription factors, the deletion of splicing factor Rbfox2 in human ESCs, where it binds to almost 7% of transcripts and autoregulates its own splicing, leads to cells death (103). These studies underscore the importance of alternative splicing in cell fate determination. From our results, the loss of KDM3A and KDM3B leads to a disruption of the pluripotency continuum and leaves the cells skewed towards the ground state (2i/LIF). If ESCs are unable to exit the pluripotent state properly, this would lead to differentiation defects and may explain the early developmental lethality observed in KDM3A/KDM3B double-knockout mice. The loss of KDM3A/KDM3B leads to the improper splicing of *Rbfox2, Tbx3, Dnmt3b,* and *Tcf12*, all of which have been implicated in the proper maintenance of pluripotency (103–106). The variant of RBFOX2 affected by the KDM3 proteins prevents higher order assembly that affects splicing regulation. As ESCs are switched from serum to 2i/LIF media, an environment of DNA hypomethylation is rapidly induced, thought to be driven by *Prdm14* and downregulation of *Dnmt3a/3b* expression (107, 108). Our results implicate another layer of DNA hypomethylation from an increase in the catalytically inactive spliced isoform of DNMT3B detected in the loss of KDM3A/KDM3B. Interestingly non-catalytic activity of DNMT3A in splicing has been implicated in embryonic stem cell differentiation (109). *Tbx3* is of particular interest because the expression level has been linked to the cell’s lineage commitment (110) and a specific alternative splicing event at exon 2 determines the protein’s binding ability to the core pluripotency regulator Nanog, controlling transcriptional output (86).

How are the splicing targets of KDM3A and KDM3B selected? One avenue could be through modulation of PRMT5 activity. PRMT5 is an arginine methyltransferase that can target both cytoplasmic and chromatin-bound proteins to modulate function (111). We examined a published PRMT5 methylome dataset and found that several splicing-associated proteins that we identified as KDM3A and KDM3B interactors such as SRSF9, SNRNP40, HNRNPA3, and SRRM2 were also methylated by PRMT5 (111) (Supplementary Figure S4F). This indicates that KDM3A/KDM3B could be serving as a scaffolding or targeting protein for PRMT5 catalytic targets. Another avenue could be by RNA modifications (112). N6-methyladenosine (m6A) regulates the fate of mRNA molecules by altering stability, localization, and splicing (113, 114). Misregulation of m6A has developmental consequences as depletion of m6A from epiblast blocks naïve pluripotency exit through the retention of methylated transcripts and results in early embryonic lethality (115). In mESC, m6A on newly transcribed RNA leads to the recruitment of KDM3B and subsequent gene activation by the removal of H3K9me2 (58). These observations imply a connection where RNA methylation precedes histone demethylation. However, we did not find a reciprocal relationship, where KDM3B splicing targets were preferentially m6a methylated (data not shown). Given the necessity of fine-tuning post-transcriptional regulation through alternative splicing for development, further study of the mechanism behind KDM3A and KDM3B splicing regulation is warranted.

Beyond pluripotency, there are examples of KDM3A affecting the splicing of single genes in cancer cells. In prostate cancer cells, KDM3A serves as a co-activator for the alterative splicing of androgen receptor variant 7, a major driver of therapy resistance in prostate cancer progression (116). In response to DNA damage in breast cancer cells, KDM3A is phosphorylated, and regulates SAT1 alternative splicing, independent of demethylating activity (117). Taken together our results emphasize this additional layer of post transcriptional gene regulation in the non-transformed developmental context and suggest that known phenotypes of the individual KDM3A and KDM3B knockouts in metabolism and germline specification will have to be re-examined.

## DATA AVAILABILITY

The datasets supporting the conclusions of this article are available in the National Center for Biotechnology Information Gene Expression Omnibus (NCBI GEO) repository with the accession code GSE230341 at https://www.ncbi.nlm.nih.gov/geo/query/acc.cgi?acc=GSE230341

The mass spectrometry proteomics data have been deposited to the ProteomeXchange Consortium via the PRIDE (118) partner repository with the dataset identifier PXD042735 and 10.6019/PXD042735. Supplementary Data are available at NAR online.

## FUNDING

This was supported by the National Institutes of Health [1R01HD105151 to R.S.]; The UW-Madison Stem Cell and Regenerative Medicine Center fellowship to C.M.D.; National Institutes of Health [5T32GM007133 to H.C.]; National Institutes of Health [R01GM141544 to P.L.B.]; National Institutes of Health [T32 GM135134 D.E.E.]; UW Genetics and Genomics distinguished undergraduate fellowship to E.C.M; Funding for open access charge: NIH [1R01HD105151].

## Conflict of interest statement

None declared.

## ACKNOWLEDGEMENTS

We would like to thank Dr. Coral Wille for ChIP-seq analysis and Hailey Thurston for technical assistance; members of the Sridharan lab for discussion and critical reading of the manuscript; and the Mass Spectrometry Core facility at UW-Madison Biotechnology Center. Figures 2A, 4B 5A, 6A, Supplementary Figure 2A, and graphical abstract created with BioRender.com. Figure 4B adapted from “mRNA Splicing Types”, by BioRender.com (2023). Retrieved from https://app.biorender.com/biorender-templates.

## Notes

### Competing Interest Statement

The authors have declared no competing interest.

### Summary of Updates

Main figures and supplemental figures updated with new data related to coIP, ChIp-qPCR, and rescue experiments. Andrew I Tak added to author list.

