## Supplemental Figures+Legends for "KDM3A and KDM3B Maintain Naïve Pluripotency Through the Regulation of Alternative Splicing"

### Supplementary Figure 1:

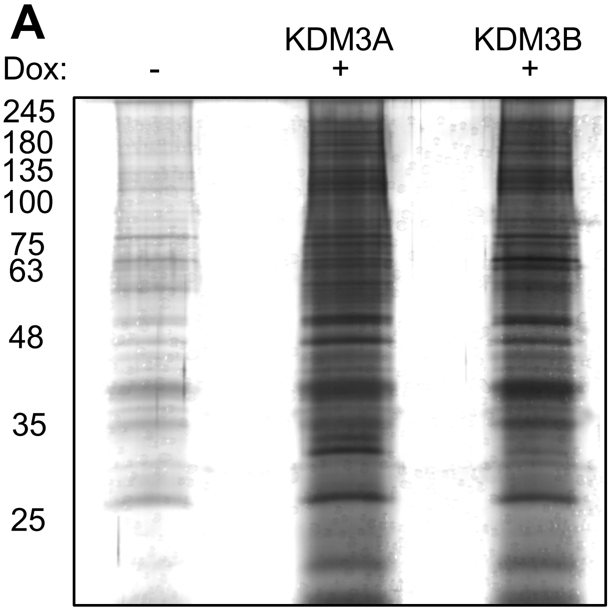


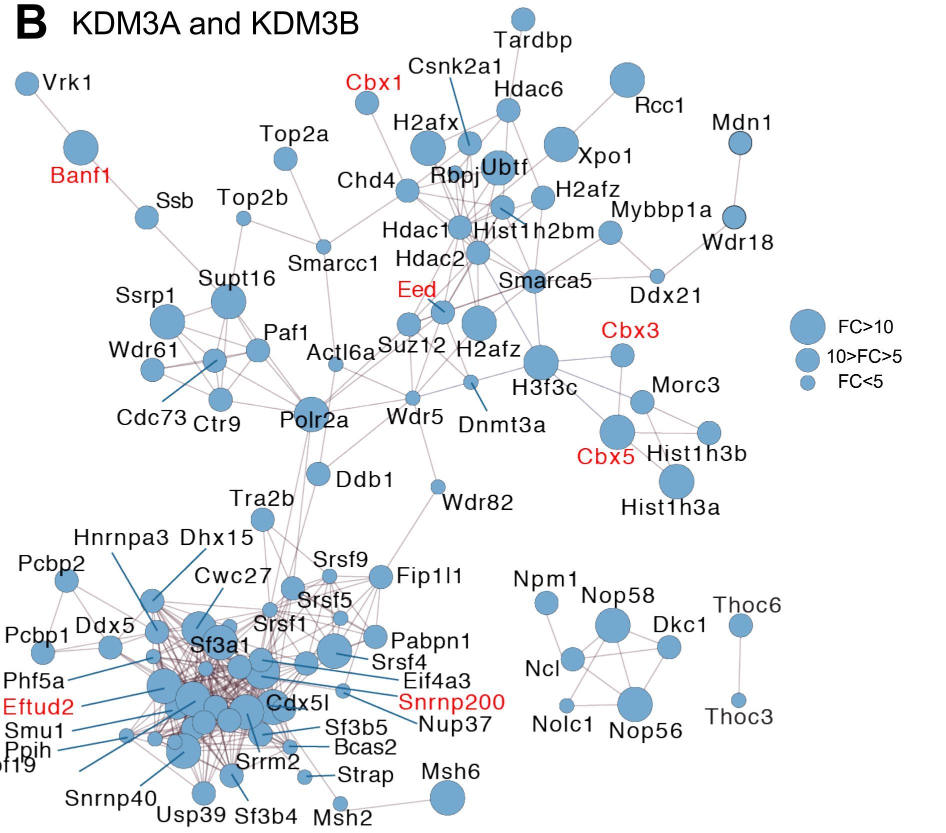


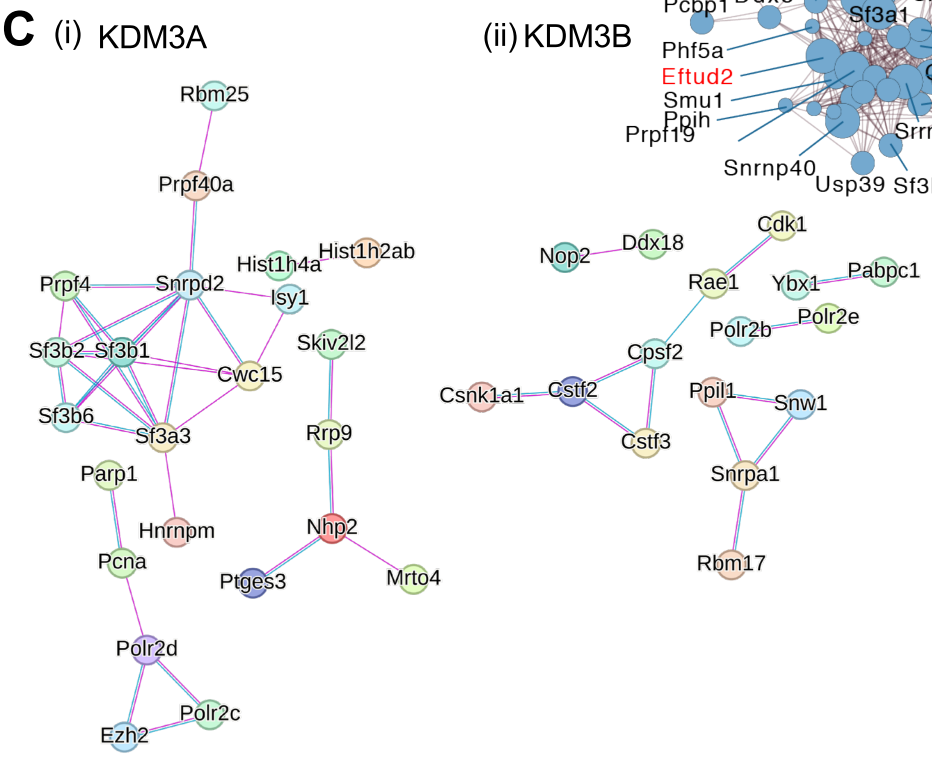


D

| Interacting Protein | Copy Number (per cell) |
|---|---|
| KDM3A | $2.8 \times 10^4$ |
| KDM3B | $1.4 \times 10^5$ |
| EFTUD2 | $1.1 \times 10^6$ |
| SNRNP200 | $7.7 \times 10^5$ |
| PRMT5 | $9.3 \times 10^5$ |

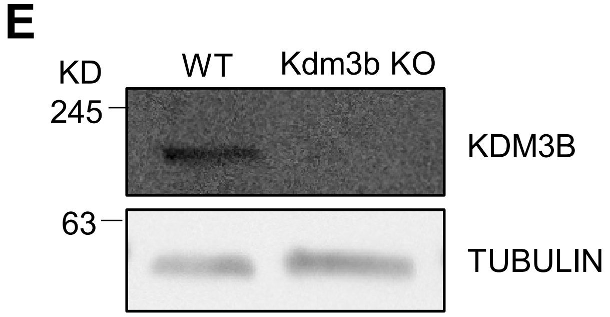


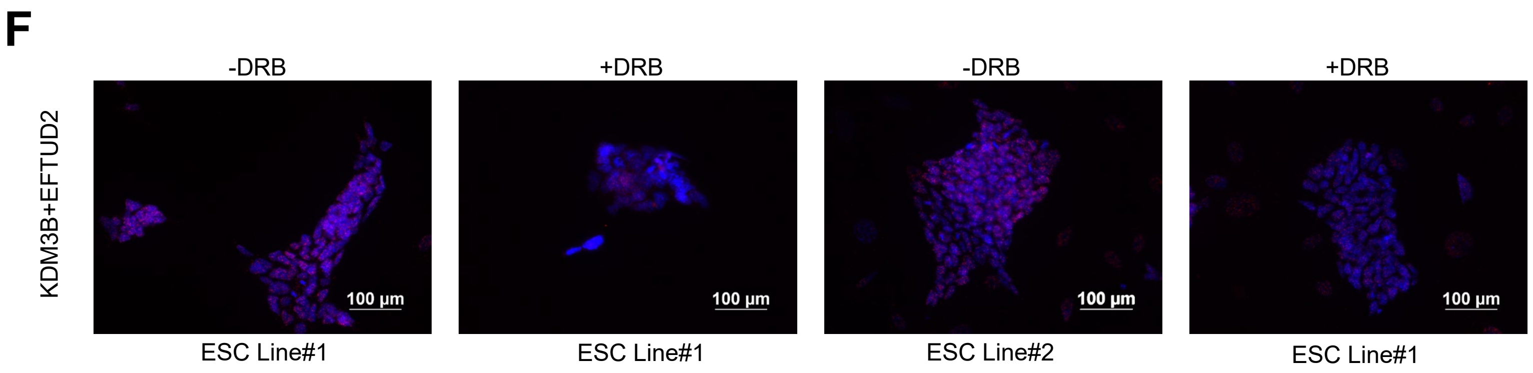

Supplementary Figure 1

(A) Silver stain of elute from FLAG-mediated immunoprecipitation from uninduced (-), FLAG-KDM3A, or FLAG-KDM3B induced ESCs.

(B) Representation of full string network depicted in Figure 1C. Example interactors from heterochromatin associated proteins, RNA splicing, and nuclear lamina associated groups are listed in red.

(C) String network of (i) KDM3A (ii) KDM3B unique interactors.

(D) Estimated number of copies of the protein per cell, derived from whole cell mass spectrometry (67) (Data obtained from https://opencell.czbiohub.org/).

(E) Immunoblot for KDM3B in WT ESCs or KDM3B knockout ESCs. Loading control = TUBULIN.

(F) Conventional fluorescence image of Proximity Ligation Assay for KDM3B and EFTUD2 with or without DRB treatment in two ESC cell lines. Scale bar = 100μM with 20x objective.

### Supplementary Figure 2:

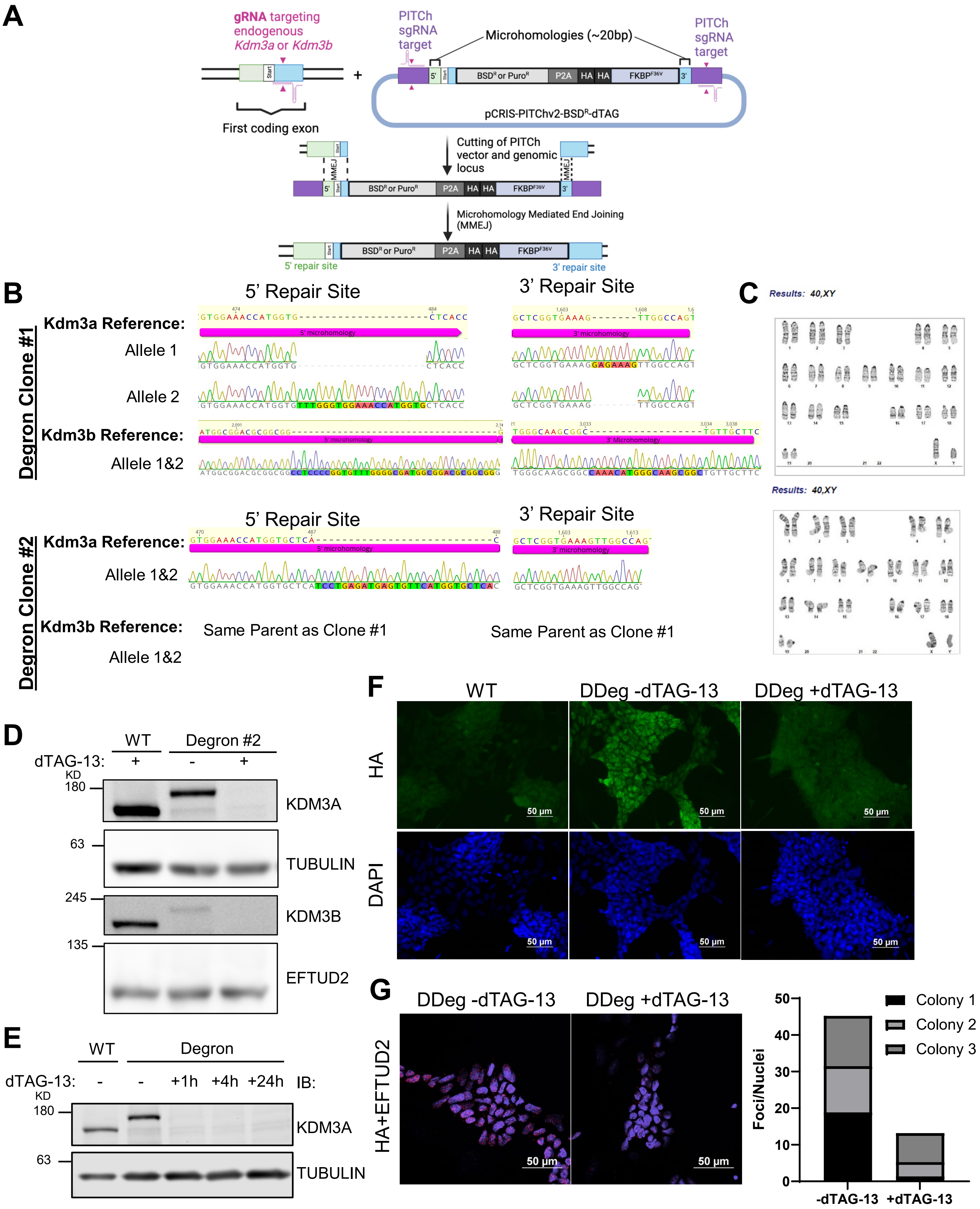

Supplementary Figure 2

(A) Schematic of PITCh (32) (Precise Integration into Target Chromosome) based CRISPR-Cas9 strategy used for inserting degron (31) tag to endogenous loci of KDM3A and KDM3B.

(B) Sanger sequencing results of 5' and 3' insertion sites of two degron clones.

(C) Normal karyotype representation of both degron clones post derivation.

(D) Immunoblot for KDM3A and TUBULIN (loading control) and KDM3B and EFTUD2 (loading control) in second degron clone. dTAG-13 treated samples harvested 24 hours after induced degradation.

(E) Immunoblot for KDM3A, KDM3B, and TUBULIN (loading control) in 24 hours timecourse of dTAG-13 induced degradation.

(F) Conventional fluorescence image of HA-staining in WT ESCs or degron ESCs +/- dTAG-13 degradation. Scale bar = 50μM with 20x objective.

(G) Single slice of 1 uM confocal images of *in situ* PLA in degron ESCs. HA+EFTUD2 antibodies in +/- dTAG-13 degradation depicted left to right. Scale bar = 50μM with 60x objective. PLA foci and nuclei were quantified in ImageJ from one colony per field of view. The ratio of foci per nuclei is reported for each colony.

### Supplementary Figure 3:

A

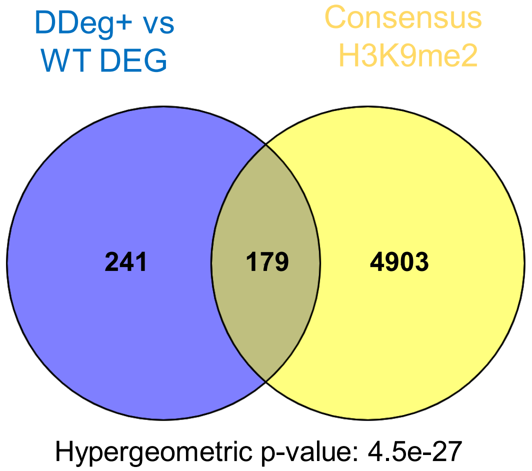


B

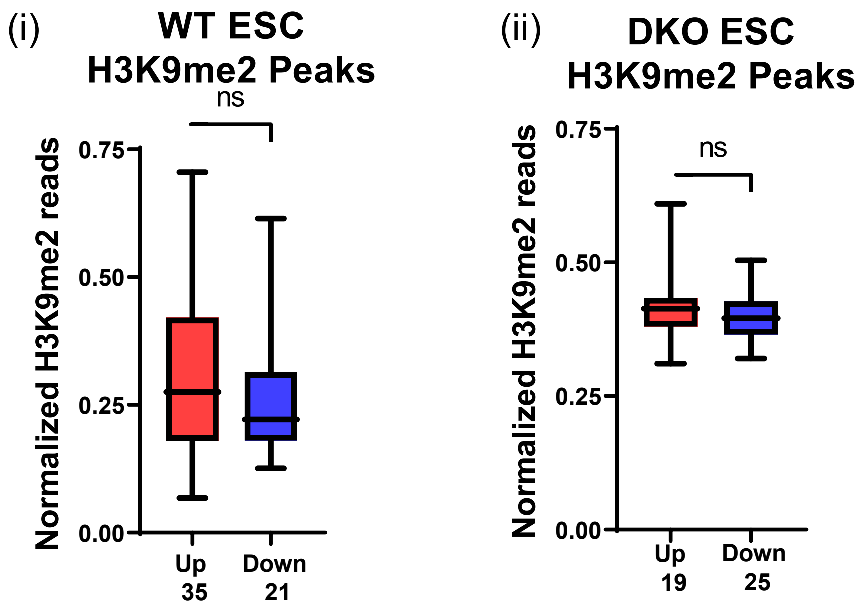

Supplementary Figure 3

(A) Two-way venn overlap of DEG and consensus H3K9me2 peaks (57, 58).

(B) Boxplot of total H3K9me2 enrichment of DEG in (i) WT ESCs or (ii) KDM3A and KDM3B DKO ESCs (22).

#### Supplemental Figure 4:

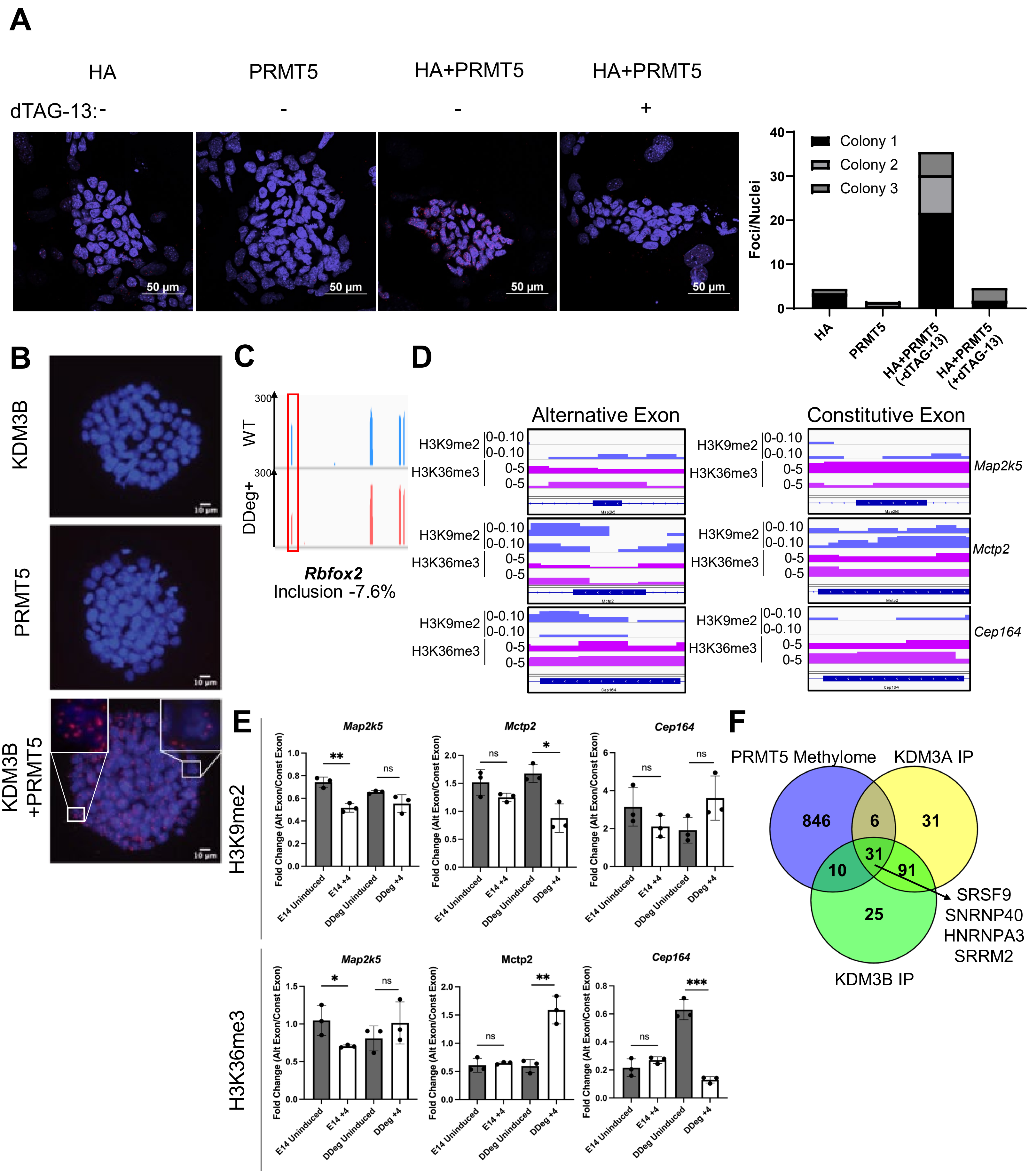

Supplementary Figure 4:

(A) Single slice of 1 uM confocal images of *in situ* PLA in degron ESCs. HA single antibody, PRMT5 single antibody, and HA+PRMT5 antibodies in +/- dTAG-13 degradation depicted left to right. Scale bar = 50μM with 60x objective. PLA foci and nuclei were quantified in ImageJ from one colony per field of view. The ratio of foci per nuclei is reported for each colony.

(B) Conventional fluorescence image of *in situ* PLA in degron ESCs. KDM3B single antibody, PRMT5 single antibody, or KDM3B+PRMT5 in second ESC line. Scale bar = 10μM with 20x objective.

(C) IGV tracks for Rbfox2 RNA sequencing. PSI value calculated in rMATs from 3 replicates in each condition. Negative value indicates event is included more in the WT condition. The affected exon is marked in red.

(D) IGV tracks from published ChIP-seq data for H3K9me2 (57) or H3K36me3 (45) at either an alternatively spliced exon called in rMATs or a constitutively spliced exon not detected in rMATs.

(E) Separate experimental ChIP-qPCR replicate of WT and inducible KDM3A/KDM3B double degron (DDeg) ESCs uninduced or treated for 4 hours with dTAG-13. Left, H3K9me2 ChIP-qPCR. Right, H3K36me3 ChIP-qPCR. Three technical replicates were performed for each condition which include alternate and constitutive exons. Fold change is determined by dividing alternative exon replicate over constitutive exon replicate.

(F) Three-way Venn diagram overlap of PRMT5 methylome (67), and KDM3A or KDM3B FLAG mediated IP results related to Figure 1.

### Supplemental Figure 5 (Related to Figure 6)

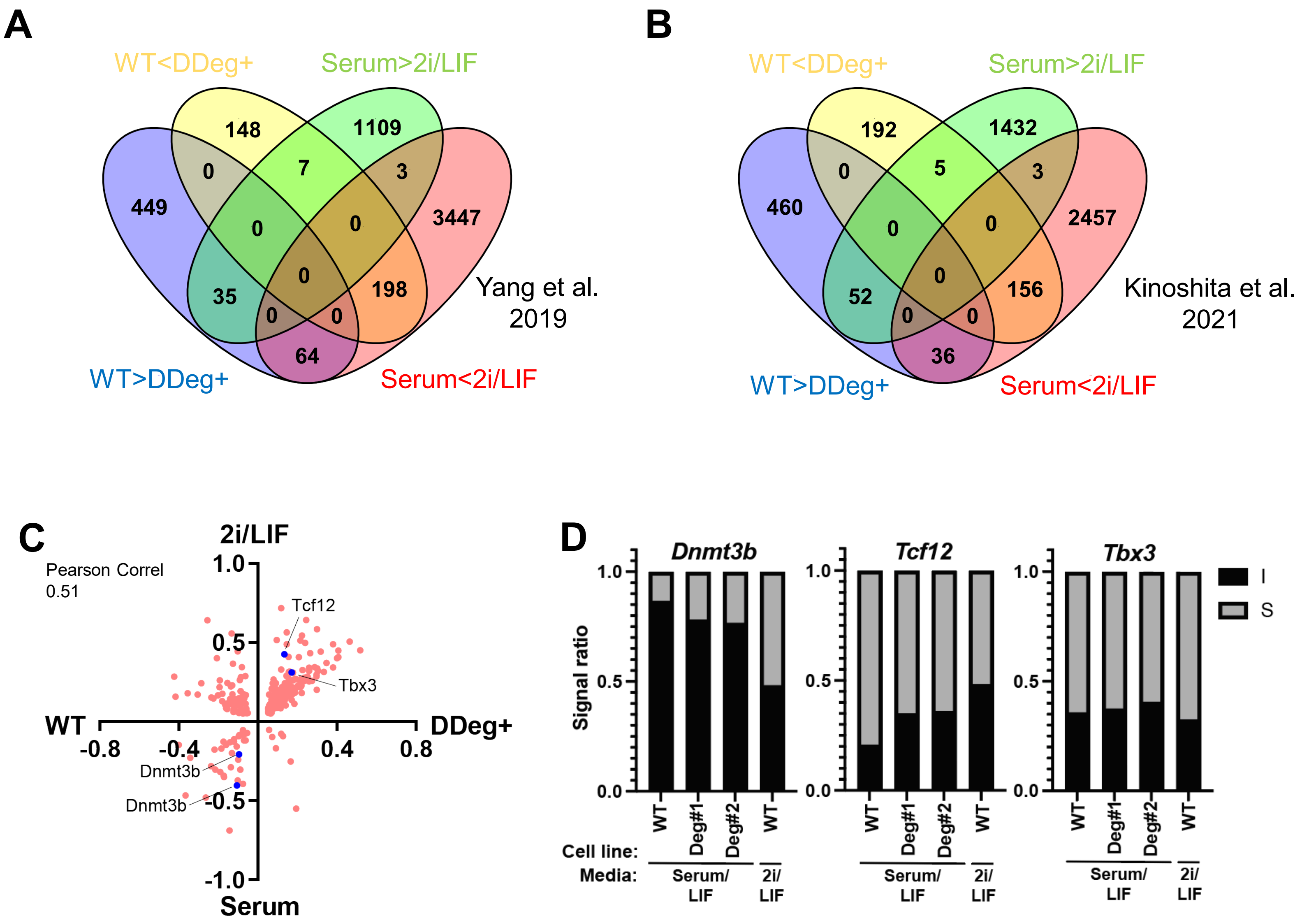

Supplementary Figure 5 (Related to Figure 6):

(A) Four-way Venn diagram overlap of aberrant splicing events called in rMATs between WT vs induced DDeg+ comparison as well as WT vs 2i/LIF (81) comparison.

(B) Four-way Venn diagram overlap of aberrant splicing events called in rMATs between WT vs induced DDeg+ comparison as well as WT vs 2i/LIF (82) comparison.

(C) Scatterplot of splicing events called by rMATs in both WT vs DDeg+ ESCs as well as serum vs 2i/LIF ESCs (82) measured by inclusion level difference score. Pearson correlation between the inclusion level difference of specific events called in both comparisons. Blue dots represent events chosen for further confirmation.

(D) Semi-quantitative PCR of splicing targets in WT ESCs, DDeg+ clones at 4 hours of degradation, and 2i/LIF ESCs. Similar to main figure 6C but presented as a inclusion/skipping ratio. Electrophoresis image was quantified using Licor software, fluorescence intensity reported as arbitrary units (a.u.) and represented in right panel. I=included isoform and S=Skipped isoform.
